# Accelerated Predictive Healthcare Analytics with Pumas, A High Performance Pharmaceutical Modeling and Simulation Platform

**DOI:** 10.1101/2020.11.28.402297

**Authors:** Chris Rackauckas, Yingbo Ma, Andreas Noack, Vaibhav Dixit, Patrick Kofod Mogensen, Chris Elrod, Mohammad Tarek, Simon Byrne, Shubham Maddhashiya, José Bayoán Santiago Calderón, Michael Hatherly, Joakim Nyberg, Jogarao V.S. Gobburu, Vijay Ivaturi

## Abstract

Pharmacometric modeling establishes causal quantitative relationships between administered dose, tissue exposures, desired and undesired effects and patient’s risk factors. These models are employed to de-risk drug development and guide precision medicine decisions. However, pharmacometric tools have not been designed to handle today’s heterogeneous big data and complex models. We set out to design a platform that facilitates domain-specific modeling and its integration with modern analytics to foster innovation and readiness in healthcare.

Pumas demonstrates estimation methodologies with dramatic performance advances. New ODE solver algorithms, such as coefficient-optimized higher order integrators and new automatic stiffness detecting algorithms which are robust to frequent discontinuities, give rise to a median 4x performance improvement across a wide range of stiff and non-stiff systems seen in pharmacometric applications. These methods combine with JIT compiler techniques, such as statically-sized optimizations and discrete sensitivity analysis via forward-mode automatic differentiation, to further enhance the accuracy and performance of the solving and parameter estimation process. We demonstrate that when all of these techniques are combined with a validated clinical trial dosing mechanism and non-compartmental analysis (NCA) suite, real applications like NLME fitting see a median 81x acceleration while retaining the same accuracy. Meanwhile in areas with less prior software optimization, like optimal experimental design, we see orders of magnitude performance enhancements over competitors. Further, Pumas combines these technical advances with several workflows that are automated and designed to boost productivity of the day-to-day user activity. Together we show a fast pharmacometric modeling framework for next-generation precision analytics.

---

Clinical pharmacologists, pharmacometricians, systems biologists and statisticians have been leveraging the latest advances in scientific computing to solve complex healthcare problems. The scientific analyses continue to evolve to be more complex as more data became available historically. However, the latest advances in computational science are not readily available to healthcare scientists.

Scientific computing and machine learning have continued to make strides in recent decade, improving core methodologies from multigrid preconditioners and neural network architectures all the way down to the basics of matrix multiplication. Surprisingly, one field which has remained consistent in both the methodologies and software used in practice is small systems of ordinary differential equations. The core software, such as the Hairer’s widely used Runge-Kutta methods (DOPRI5, DOP853) [13] and the backwards differentiation formulae (BDF) methods derived from LSODE [16] (LSODA [15], VODE [8], and CVODE [17]), were fully developed by the 90’s and have been the stable core of scientific computing ever since. However, in this manuscript we will demonstrate that this general area of computational science can see dramatic computational performance improvements by developing a new set of solver algorithms specialized for simulation of small-scale stiff and non-stiff dynamical systems.

The delay in the availability of the latest advances to healthcare scientists limits their ability to gain deep insights into why some patients do not respond to treatment, why some develop serious toxicity, risk factors for deciding on the right treatment for the right patient (precision medicine). The access to heterogenous data from laboratory measurements, radiographic scans, clinical scripts, genomic and genetic data has not fully translated into actionable science due to the lack of more efficient tools, to a large extent [12]. A related challenge has also has been the lack of a unified platform to perform these advanced scientific analyses. For example, a systems biologist cannot perform nonlinear mixed effects modeling for the same project within one software. Unknowingly, this caused serious communication issues between scientists at different stages of the same project. On other hand, one scientist cannot be expected to master all software tools. An integrated platform that allows scientists to build tools in a seamless manner is urgently needed.

Fields which require high fidelity and stable estimation of parameters of such dynamical systems, such as pharmacometrics and systems biology, are frequently constrained by the calculation times required when solving large numbers of such systems [2, 7]. In this manuscript we will demonstrate how these new differential equation solvers are integrated with automatic differentiation and parameter fitting routines in a manner that decreases the time of real-world applications by an order of magnitude.

Likewise, without a bridge that connects these new solver and compiler tools to domain-specific tooling, the ability for pharmacometricians to make use of these tools is limited and requires a modeler to go to tools usually written in low level languages like C++ or Fortran. This limits the flexibility of the allowed models and decreases the speed at which the latest advancements in high performance computing become available to the practitioner. Pumas (**P**harmace**U**tical **M**odeling **A**nd **S**imulation) is impacting this flow by directly packaging the latest mathematical and hardware advances inside of a pharmacometric modeling context inside of the Julia high level high performance language [6]. The driving paradigm of Pumas is to have a completely flexible core while successively simplifying interfaces through generally useful defaults. This allows for a graded approach to learning the modeling framework where beginners can simply use the defaults and expect it to match standard behavior, while keeping non-standard pharmacometric models (such as stochastic differential equations) directly accessible, optimized, and able to utilize all of the hardware compatibility tooling in a first-class manner. This is made possible because Pumas is written for the Julia programming language while being written entirely written with the Julia programming language; allowing user-written extensions to flow directly from standard usage. While Julia is a high level programming language which allows Pumas to have ease of use for non-programmers, the Julia language is a just-in-time (JIT) compiled language as fast as low level languages like C or Fortran. Thus both the library and any user-written components are free from interpreter overhead imposed in languages like Python or R. Therefore, our approach with Pumas is to not shy away from using the language and its extensive package ecosystem, and instead integrate our approaches with these tools.

In this paper we will describe the generalized nonlinear mixed effects models (NLME) [7] framework which Pumas utilizes for personalized precision dosing [31]. We will then showcase how the deep integration with the differentialEquations.jl [33] software package can allow for many domain-optimized approaches to be accessible within the context of pharmacometric models such as those seen in pharmacokinetics and pharmacodynamics (PK/PD) [3, 46]. We will demonstrate how this connection facilitates Integrated Pharmacometrics and Systems Pharmacology (iPSP) [44] by allowing the optimized solution of large sparse PBPK and QSP models within the NLME context, and showcase how alternative differential equation forms like differential-Algebraic Equations (DAEs) can be used to stabilize a model or Stochastic differential Equations (SDEs) can be used to generalize a model to include process noise. Features of Pumas, like integrated high-performance noncompartmental analysis (NCA), automatically parallelized visual predictive checks (VPCs), and fast tooling for optimal design of clinical experiments is all integrated into the Pumas system to allow full applications to be simple and fast. After seeing the modeling benefits of such a framework, we detail the performance benefits, showcasing acceleration over previous software in the standard ODE NLME cases while demonstrating automated parallelism. Together Pumas is a tool built for the next-generation of pharmacometric analysis that will allow for modeling and developing personalized precision medicine in areas that were previously inaccessible due to excessive computational cost.

## 1 Introduction to Nonlinear Mixed effects Modeling with Pumas

### 1.1 Nonlinear Mixed effects Models

Many pharmacometric studies fall into a class of models known as nonlinear mixed effects models [7]. Figure 1 gives a diagramatic overview of this two-stage hierarchical model. In the context of pharmacometrics, the lower level model describes the drug dynamics within a subject via a differential equation while the higher level model describes how the dynamical model is different between and within subject. The fixed effects are the values *θ* which are independent of the subject. One can think of the fixed effects as population typical values. With every subject *i* there is a set of covariate values *Z*_*i*_ which we can know about a subject in advance, such as their weight, height, or sex. We then allow a parameter *η*_*i*_ which is known as the random effect to be the difference between the typical value and the subject. The structural model *g* is the function that collates these values into the dynamical parameters for a subject *p*_*i*_, i.e.:

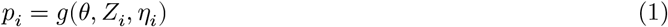

**Figure 1:**
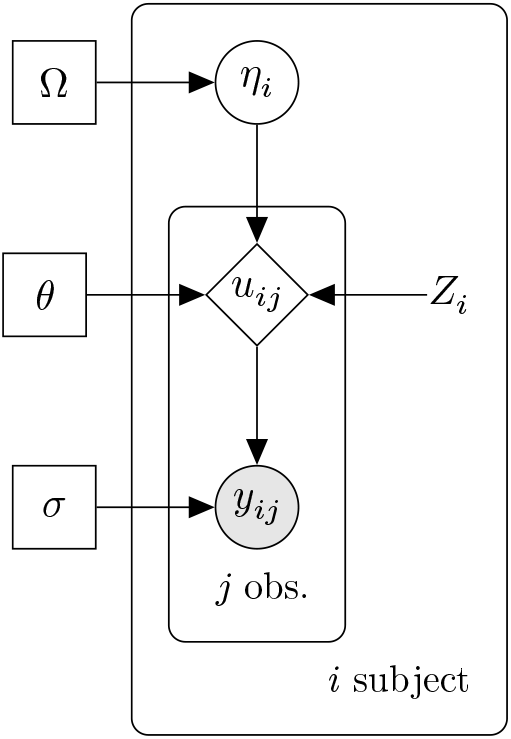
Diagrammatic description of nonlinear mixed effect models. A plate diagram of the model: rectangle nodes denote parameters, circles denote random quantities which are either latent (unfilled) or observed (filled), diamonds are deterministic given the inputs, and nodes without a border are constant.

The dynamical parameters are values such as reaction rates, drug clearance, and plasma volume which describe how the drug and patient reaction evolves over time through an ordinary differential equation (ODE):

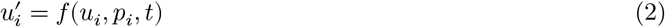

where *f* is the dynamical model and *u* is the state variables that are being evolved, such as the drug concentrations over time. The *j*th observables of patient *i, y*_*ij*_, such as the maximum concentration or area under the curve (AUC), are derived values from the dynamical simulation through a function *h*:

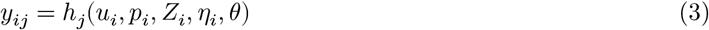

Lastly, measurements are taken on the derived values by assuming measurement noise of some distribution (commonly normal) around the prediction point.

The following showcases a classic model of Theophylline dynamics via a 1-compartment model implemented in Pumas, where patients have covariates:

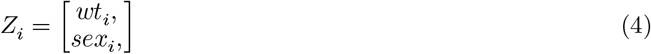

a structural collocation:

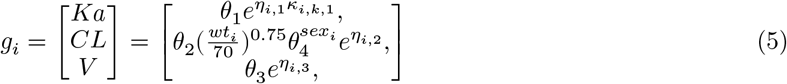

internal dynamics:

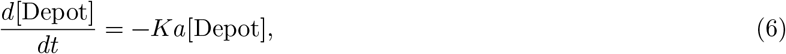

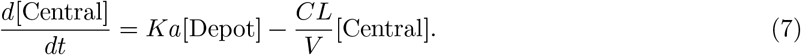

and normally distributed measurement noise. The reason for the NLME model is that, if we have learned the population typical values, *θ*, then when a new patient comes to the clinic we can guess how they are different from the typical value by knowing their covariates *Z*_*i*_ (with the random effect *η*_*i*_ = 0). We can simulate between-subject variability not captured by our model by sampling *η*_*i*_ from some representative distributions of the *η*_*i*_ from our dataset (usually denoted *η*_*i*_ ~ *N*(0, Ω)). Therefore this gives a methodology for understanding and predicting drug response from easily measurable information.

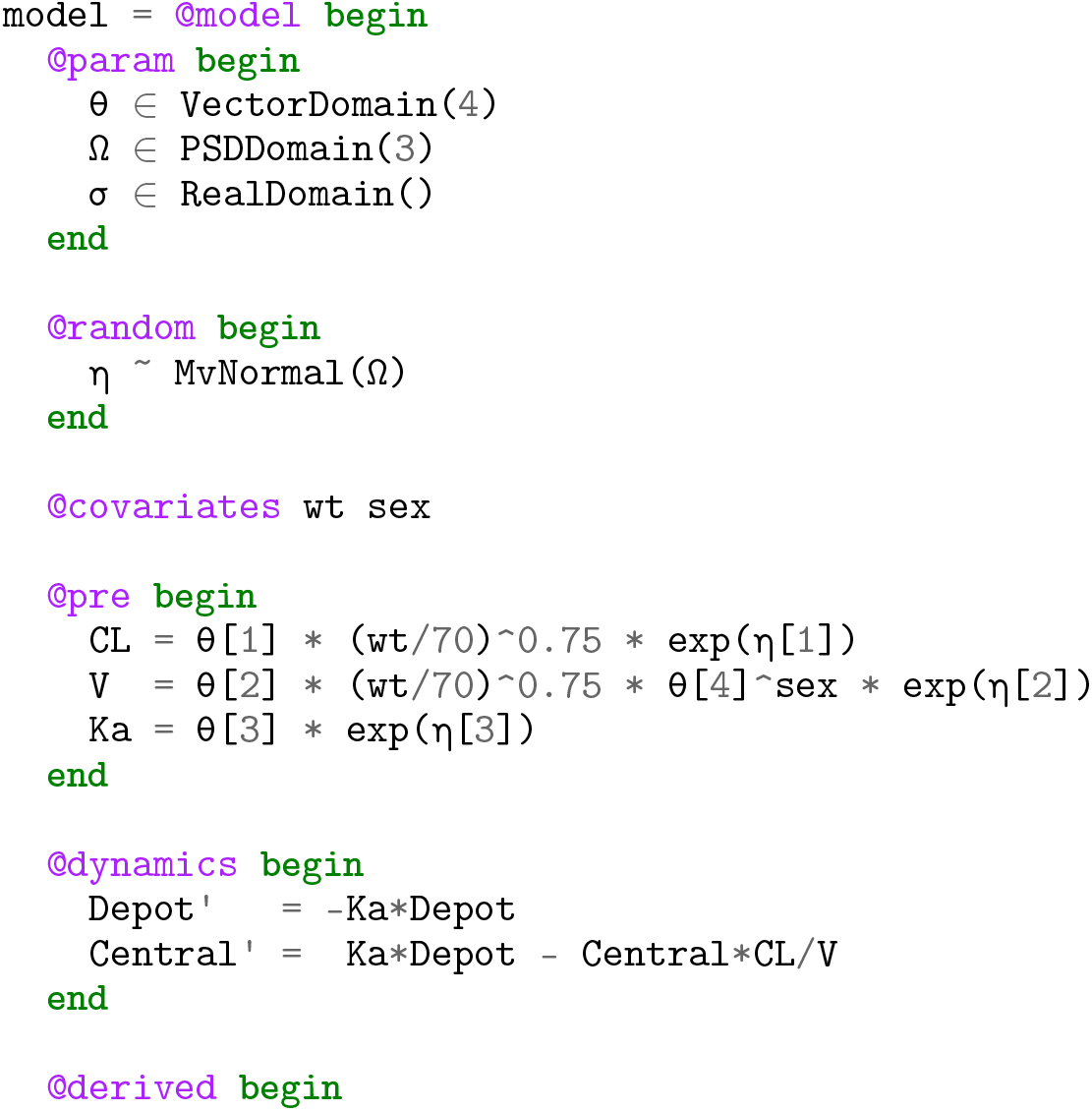

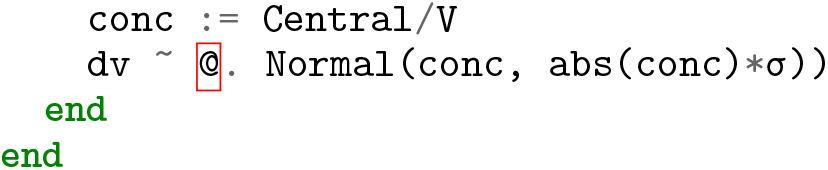

### 1.2 Clinical Dosage Regimen Modeling

In addition to nonlinear mixed effects model designations, Pumas allows for the specification of clinical dosage regimens. These dosage regimens are modeled as discontinuities to the differential equation and can be specified using a standard clinical dataset or by using the programmatic DosageRegimen method directly from Julia code. Figure 2 showcases the result of a Pumas simulation of this model with steady state dosing against the pharmaceutical software NONMEM [4], showcasing that Pumas recovers the same values. The test suite for the dosing mechanism is described in Appendix 5. Appendix Figures 11 and 12 demonstrate similar results across a 20 different dosage regimens using numerical ODE solving and analytical approach for accelerated handling of linear dynamics. One interesting feature of Pumas demonstrated here is the ability to specify the continuity of the observation behavior. Because of the discontinuities at the dosing times, the drug concentration at the time of a dose is not unique and one must choose the right continuous or the left continuous value. Previous pharmaceutical modeling software, such as NONMEM, utilize right-continuous values for most discontinuities but left-continuous values in the case of steady state dosing. To simplify this effect for the user, Pumas defaults to using the right-continuous values, but allows the user to change to left continuity through the continuity keyword argument. By using right continuity, the observation is always the one captured post-dose. Since the simulations usually occur after the dose, this makes the observation series continuous with less outlier points, a feature we believe will reduce the number of bugs in fitting due to misspecified models.

**Figure 2:**
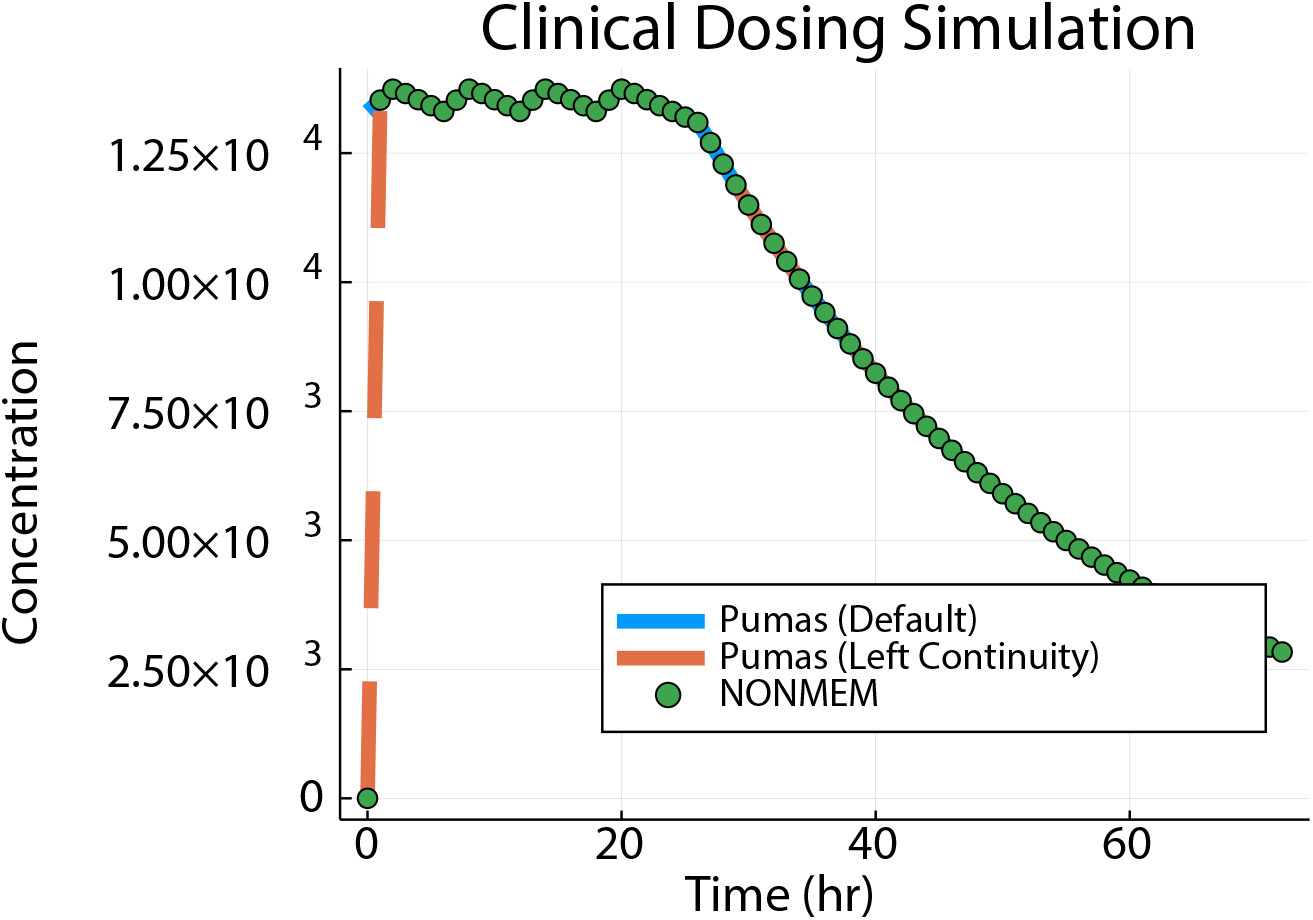
Clinical dosing simulation comparison against NONMEM. Shown is the continuous simulation trajectory outputted from Pumas using the default (right) and left continuity choices, compared against NONMEM. This figure both showcases a relative difference < 1 × 10^−10^ except at the starting point, where the NONMEM simulation requires a pre-dose estimate of zero whereas the Pumas simulations allows a continuous output via a post-dose observation and allows for switching to the NONMEM behavior.

**Figure 3:**
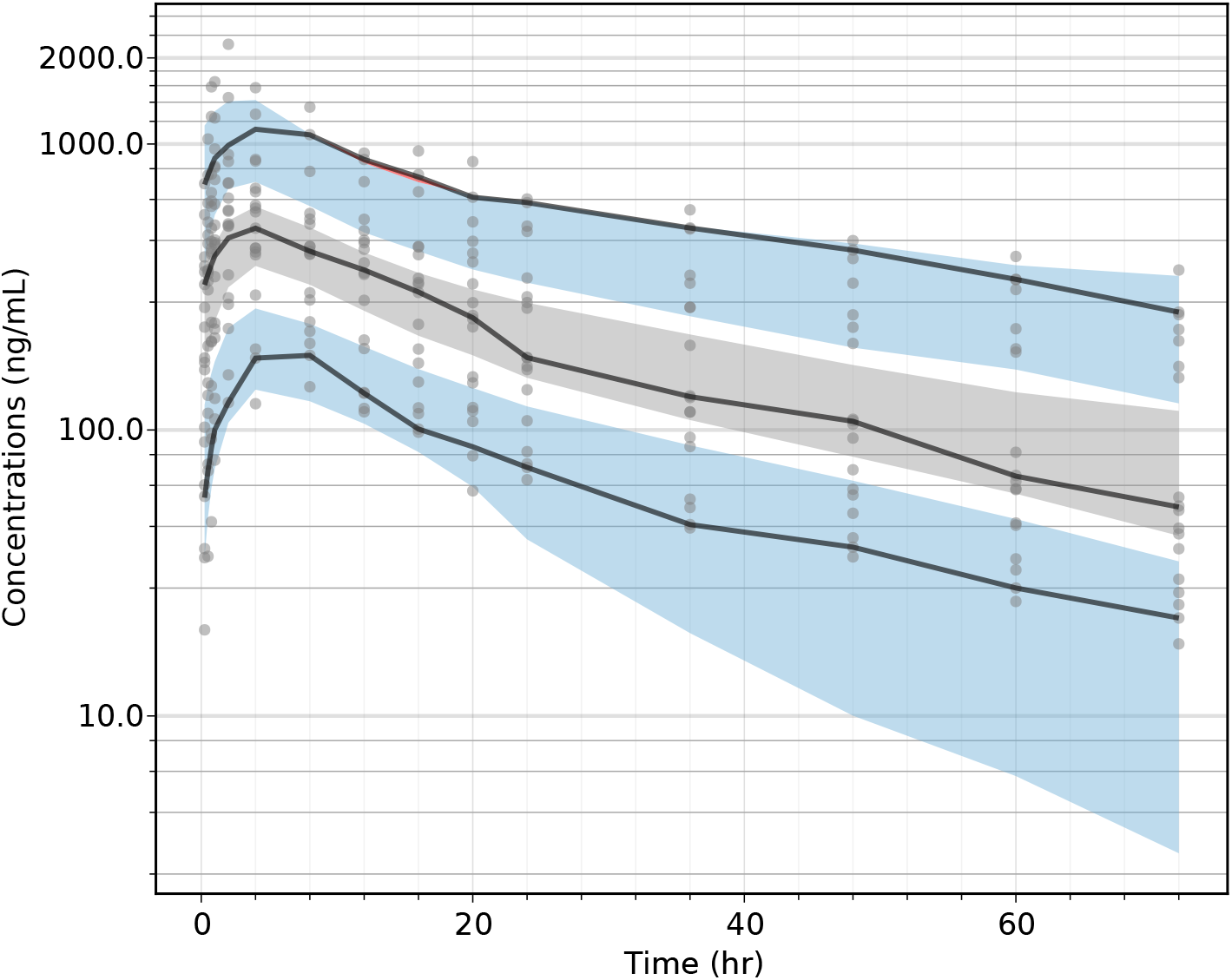
Visual Predictive Check.

**Figure 4:**
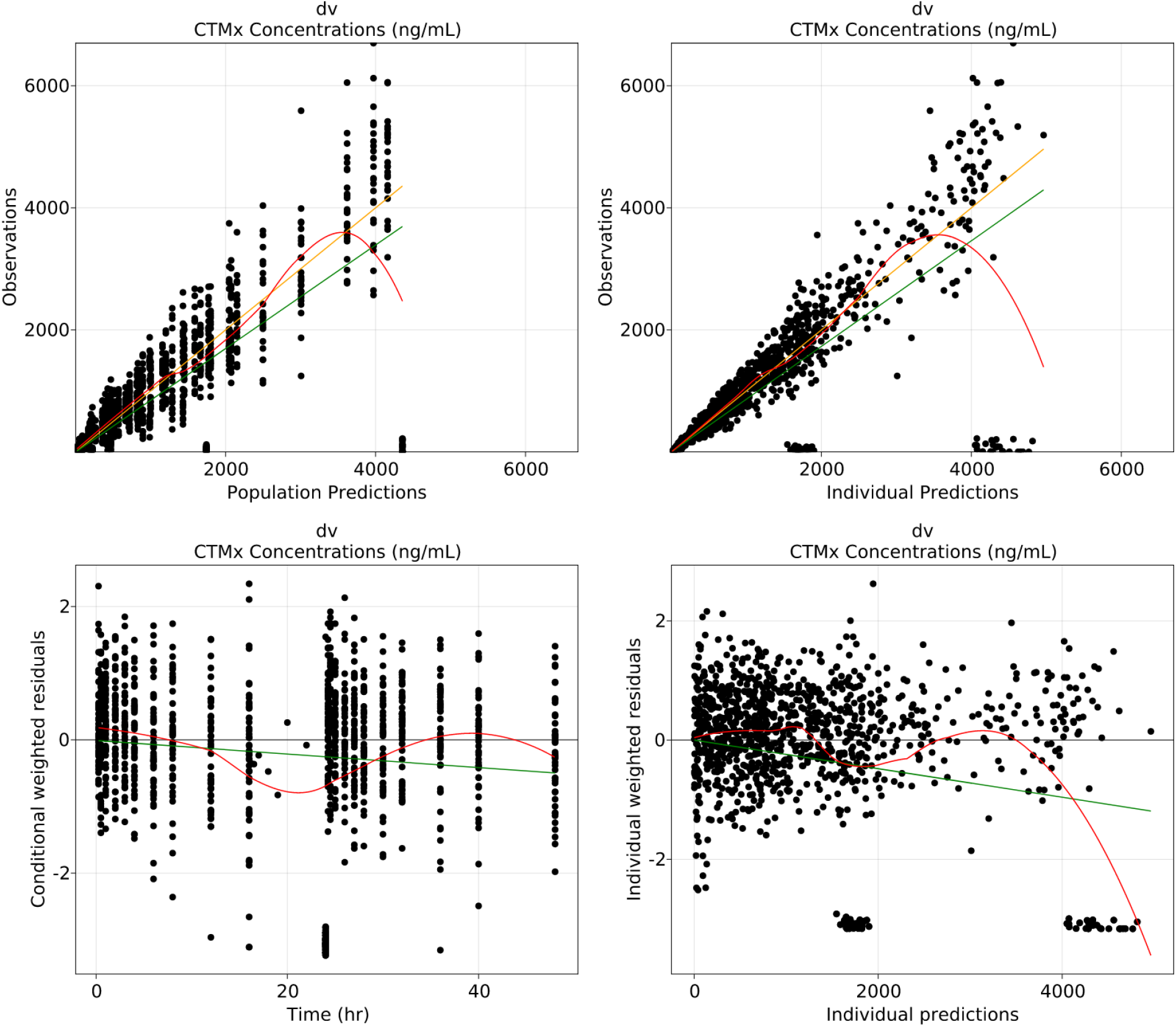
Goodness-of-fit plots for fitted model.

**Figure 5:**
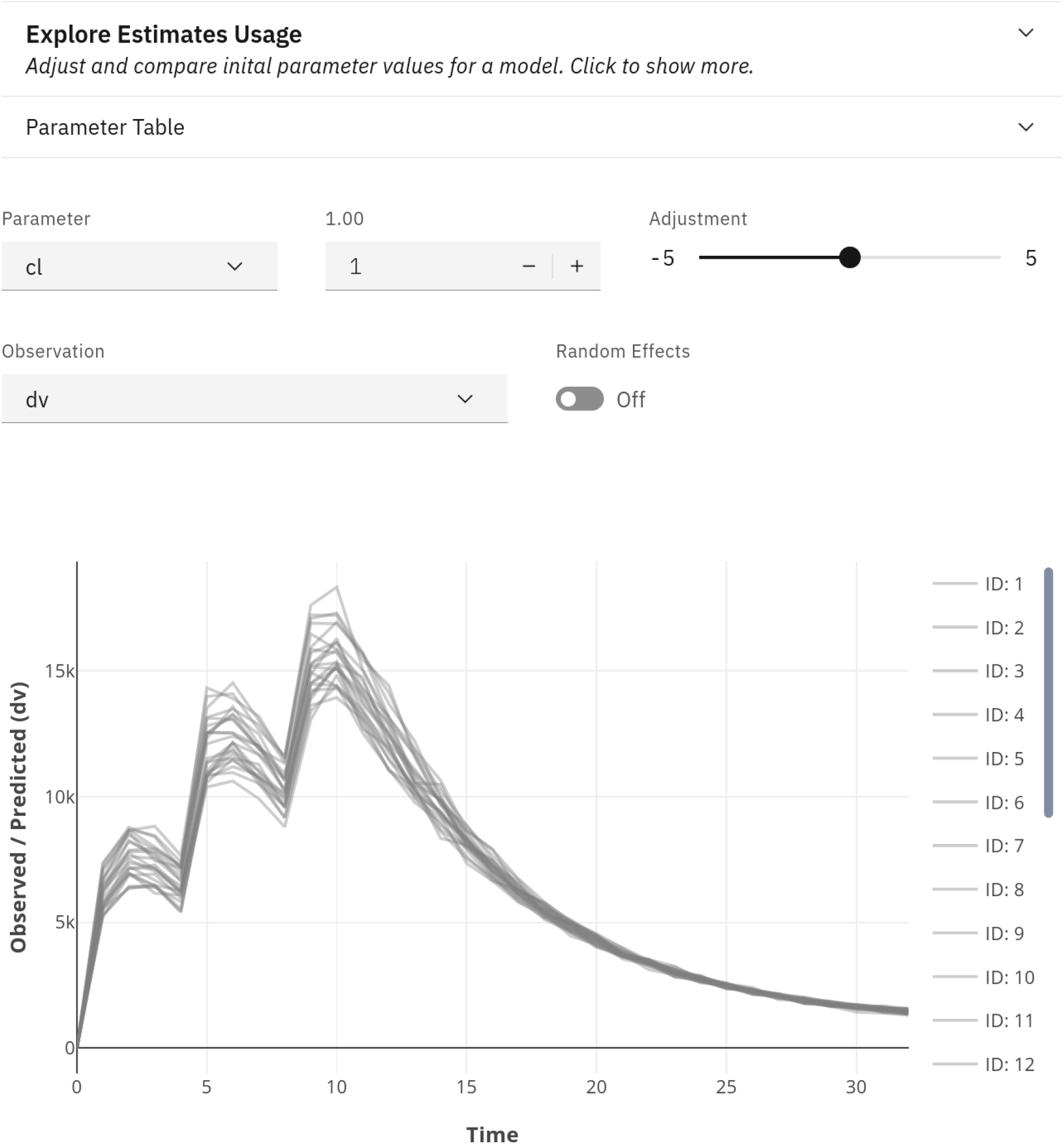
The explore estimates app. This can be used to pick initial estimates for model fitting visually.

**Figure 6:**
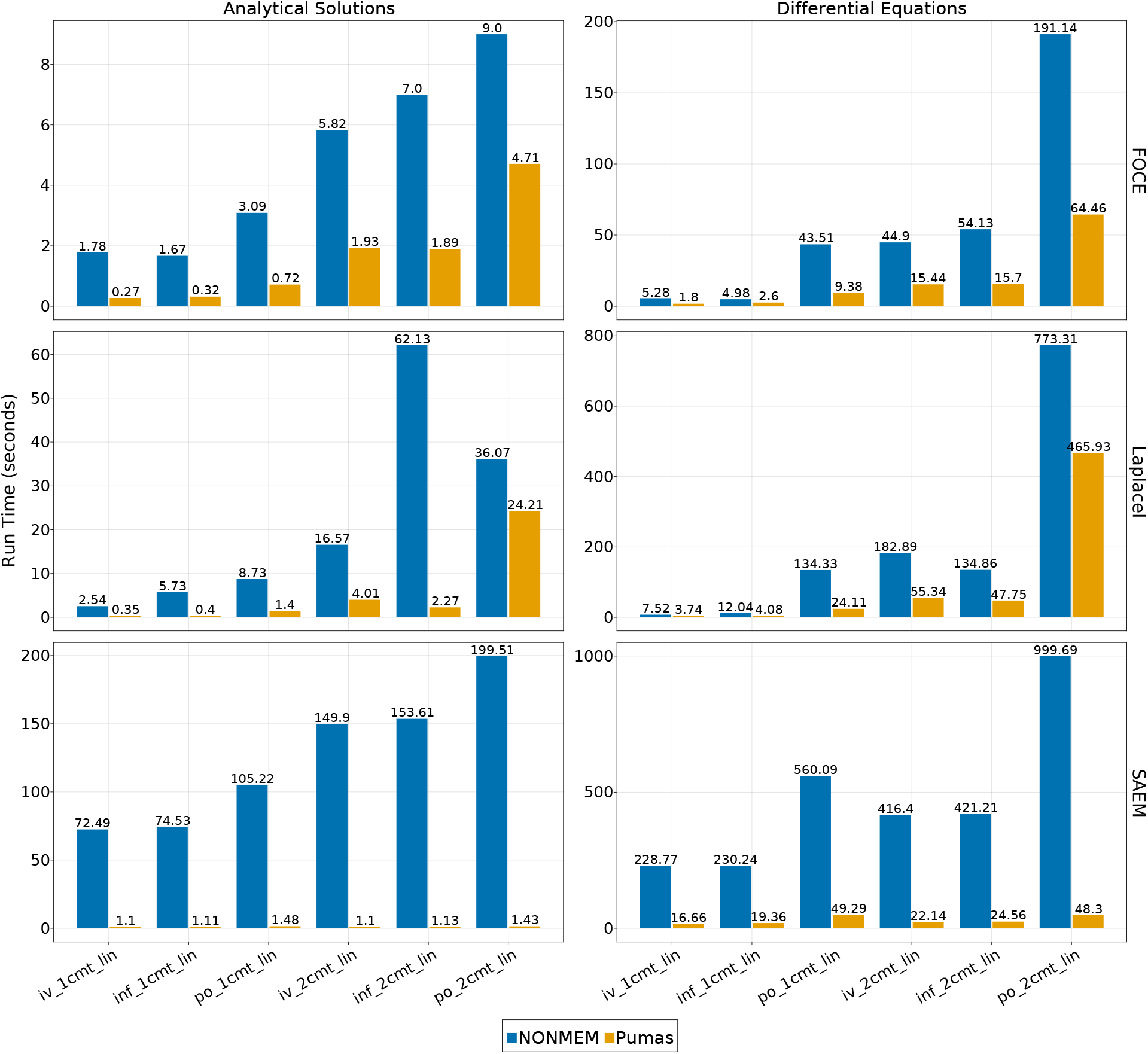
Maximum likelihood estimation and SAEM benchmark results for single dosing simulations.. Shown is the run time of a fit of the various PK models with the FOCE, LaplaceI, and SAEM population integral approximation schemes in NONMEM and Pumas. Analytical solutions refers to direct usage of the solution to the differential equation in the fitting process. For numerical approximations, NON-MEM was run using ADVAN6 (the DVERK 6th order Runge-Kutta method) while Pumas used the Vern7 ODE solver. In all these cases, there is 100 percent agreement between the final loglikelihoods and the parameter estimates between Pumas and NONMEM

**Figure 7:**
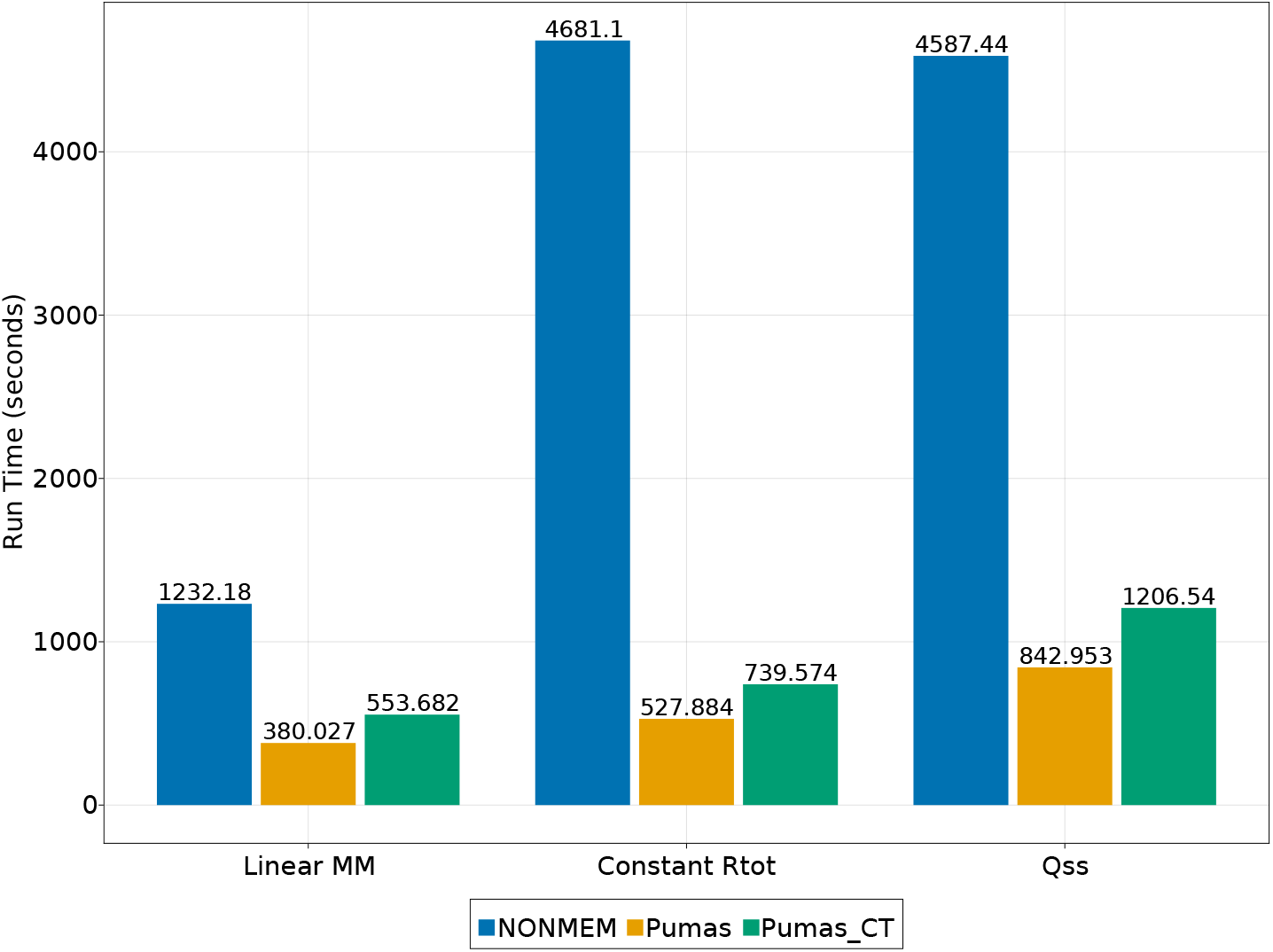
Runtime comparisons of non-linear mechanistic TMDD models. Shown is the run time of a fit of the Michaelis-Menten (MM) model, Constant Rtot model and Rapid binding (QE) and quasi steady-state (QSS) models. Shown are the NONMEM runtime, the Pumas runtime, and the Pumas runtime which includes compile time. NONMEM was run using ADVAN13 TOL6 (LSODA) while Pumas used the default ODE solver for parameter estimation (Auto-switch Vern7 mixed with Rodas5). In all these cases, there is 100 percent agreement between the final loglikelihoods and the parameter estimates between Pumas and NONMEM

**Figure 8:**
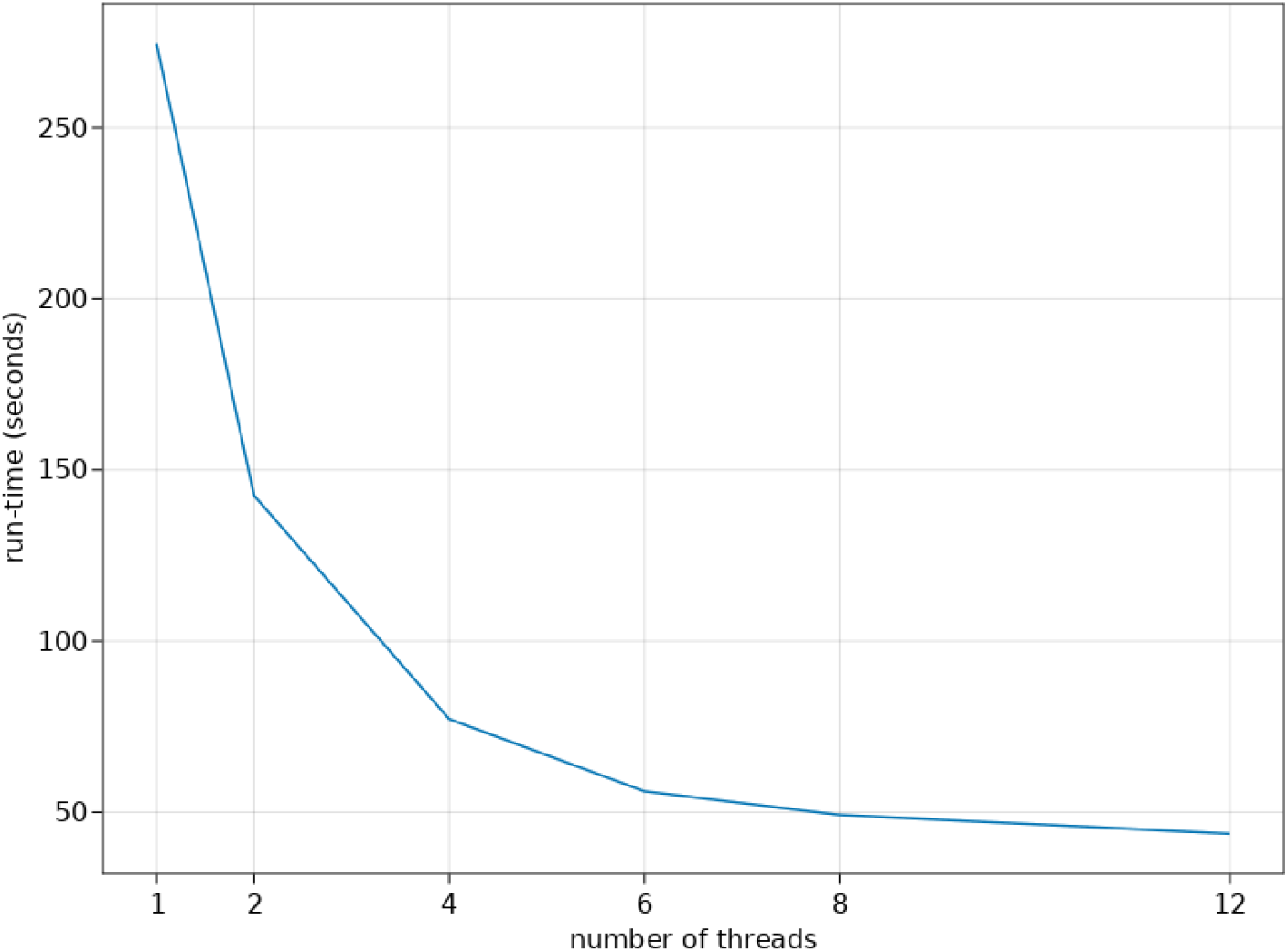
Scaling of multithreaded fitting of a HCV model. Shown is the run time of a fit of a simultaneous PK/PD model HCV model described in 5 with multithreading with 1, 2, 3, 4, 6, 8 and 12 threads.

**Figure 9:**
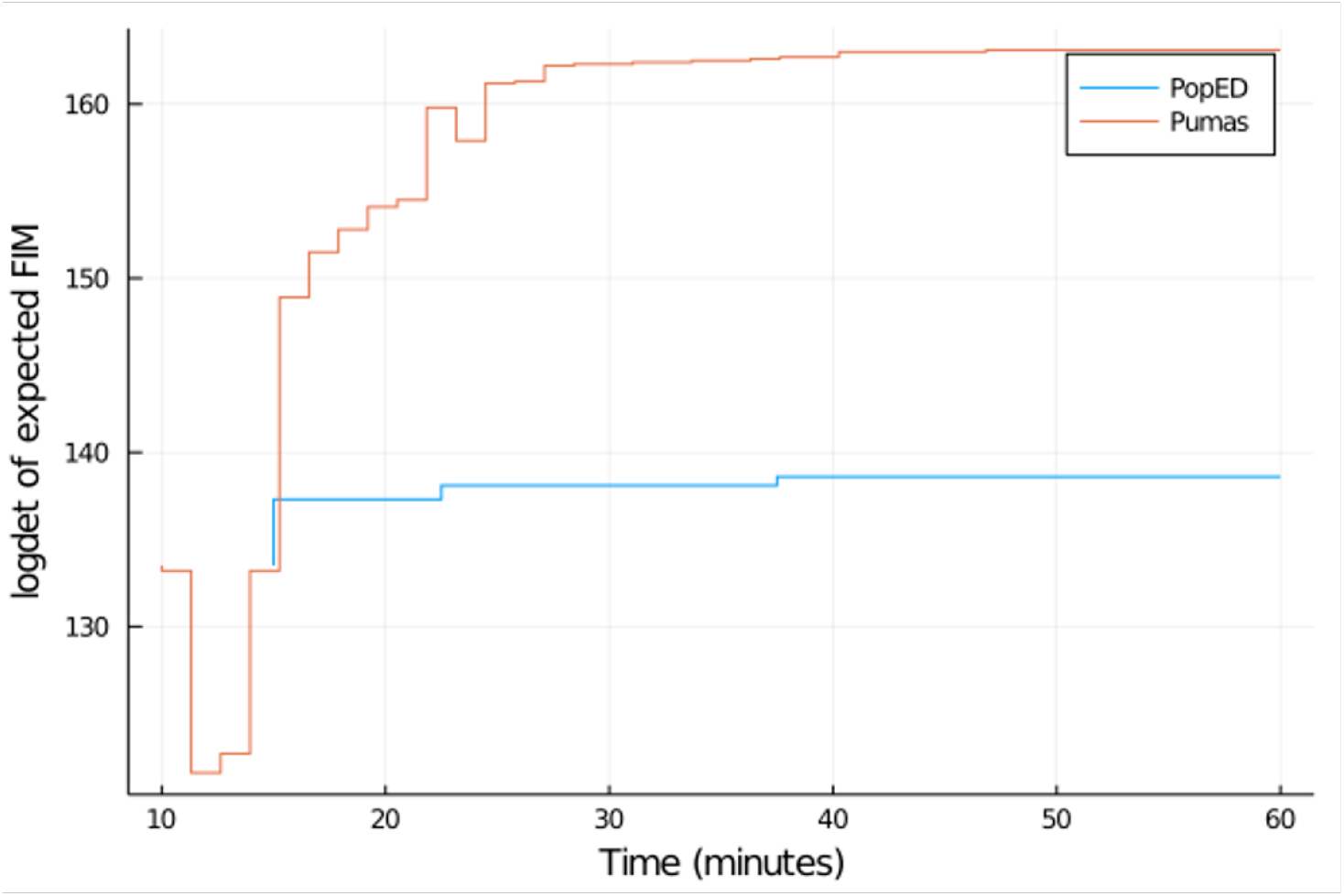
Objective value (log determinant of expected FIM) profiles for PopED vs Pumas over 1 hour of optimization.

**Figure 10:**
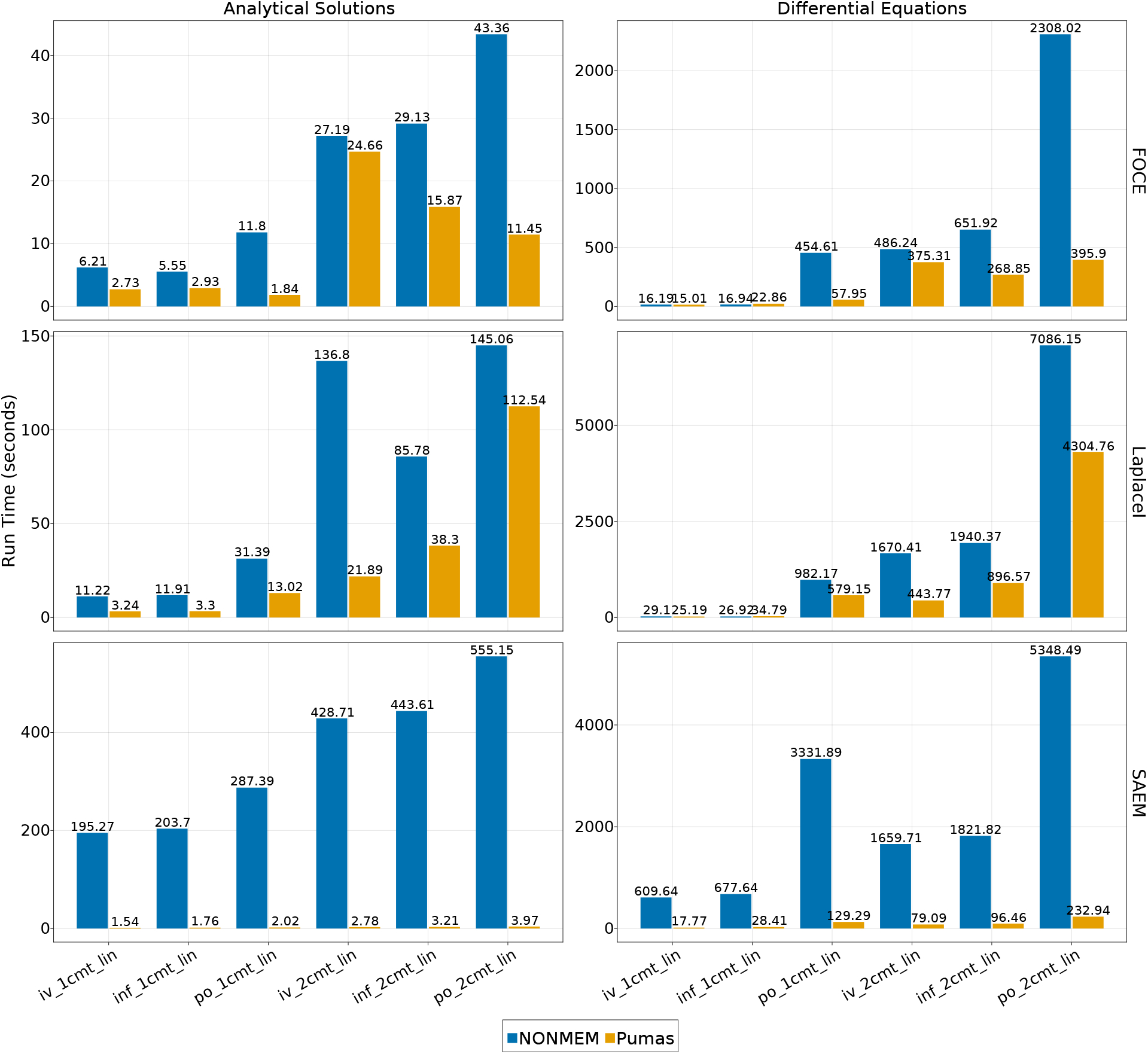
Maximum likelihood estimation and SAEM benchmark results for multiple dosing simulations.. Shown is the run time of a fit of the various Pk/Pd models with the FOCE, LaplaceI, and SAEM population integral approximation schemes in NONMEM and Pumas. Analytical solutions refers to direct usage of the solution to the differential equation in the fitting process. For numerical approximations, NONMEM was run using ADVAN6 (the DVERK 6th order Runge-Kutta method) while Pumas used the Vern7 ODE solver.

**Figure 11:**
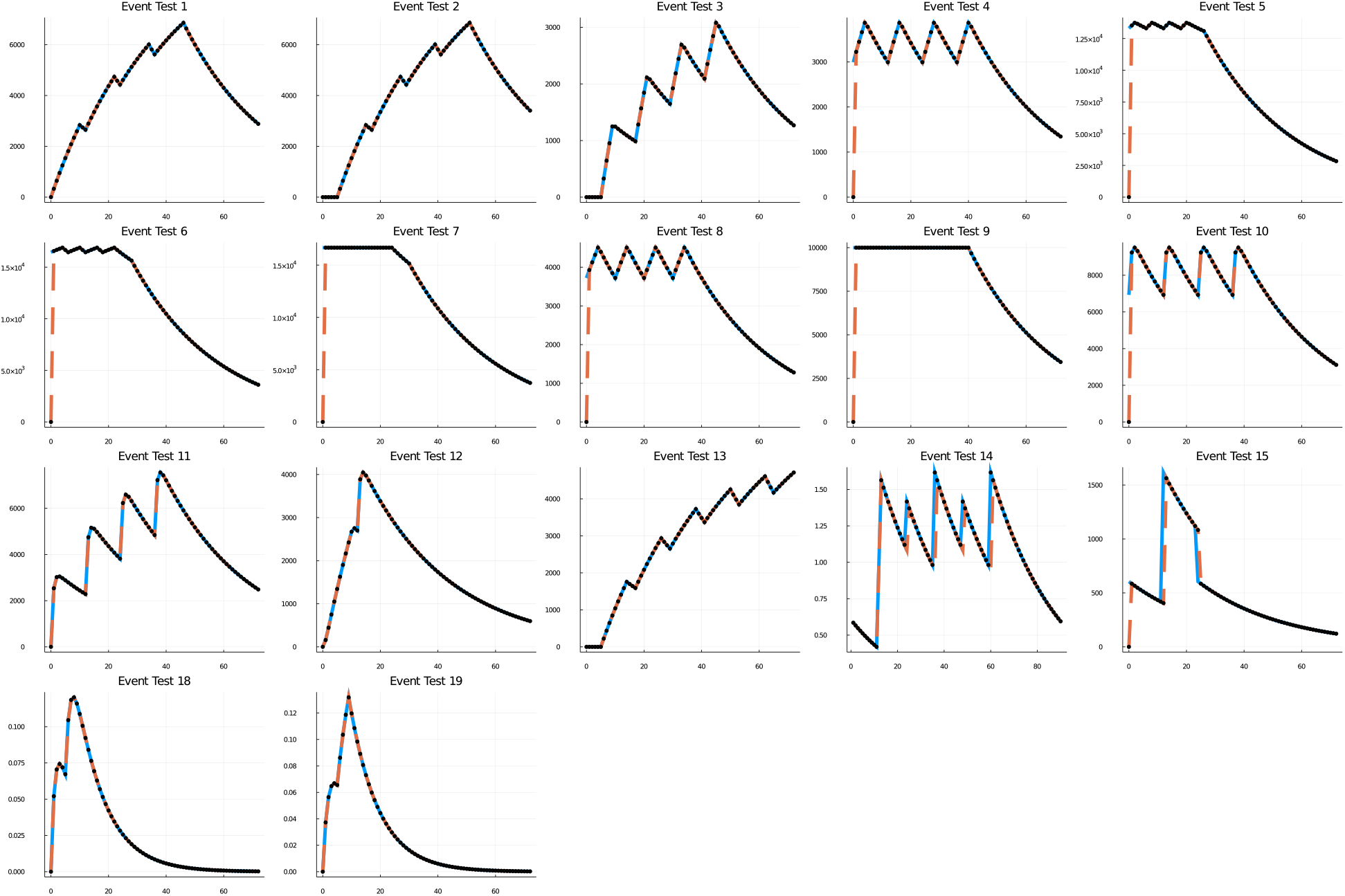
Clinical dosing simulation verification against NONMEM for analytical solutions. Shown are the clinical dosing simulation verification of the models described in Section 5 using analytical solutions of the differential equation mixed with the event handling system.

**Figure 12:**
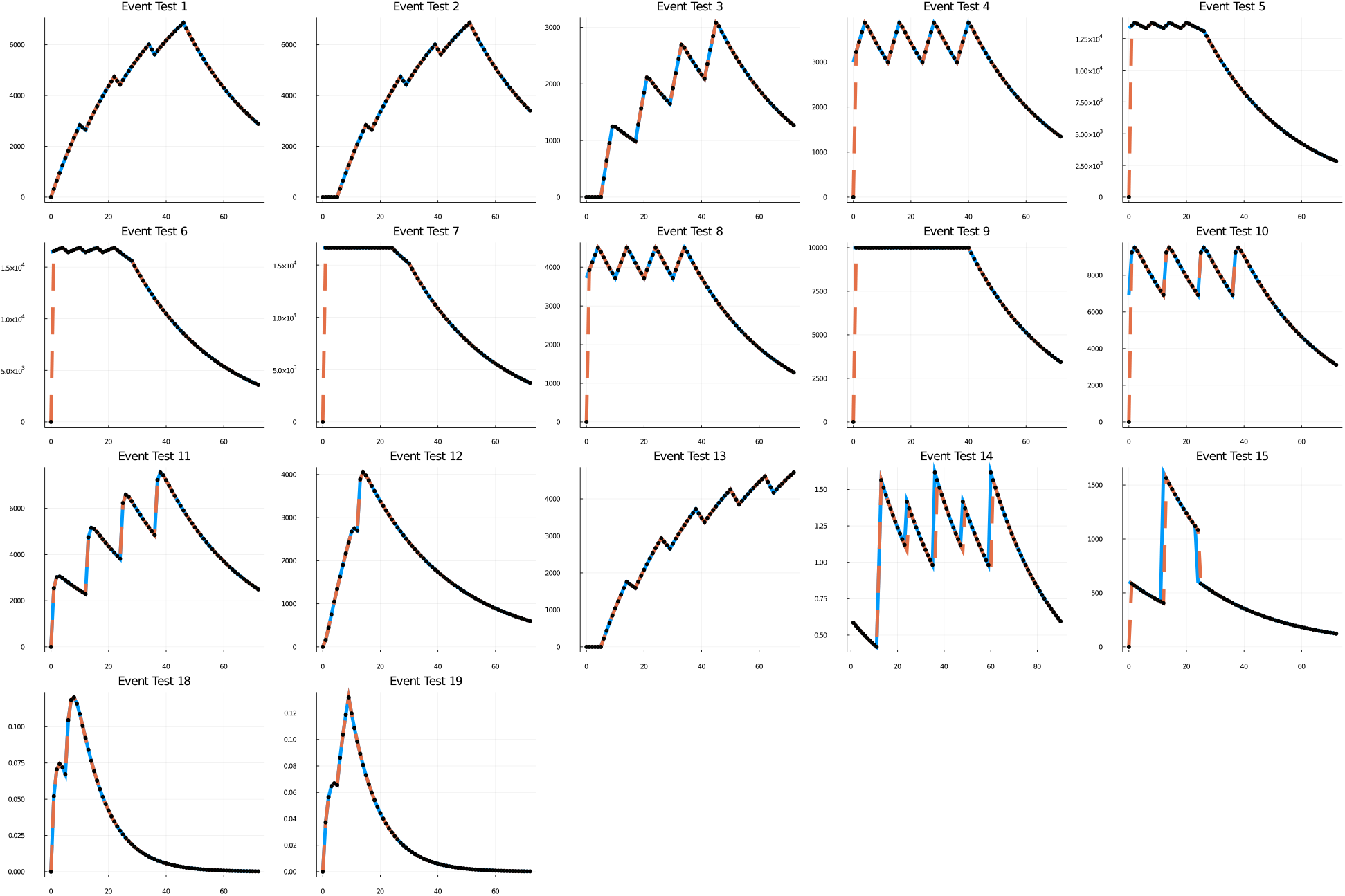
Clinical dosing simulation verification against NONMEM for analytical solutions. Shown are the clinical dosing simulation verification of the models described in Section 5 using numerical approximations of the solutions of the differential equation mixed with the event handling system.

### 1.3 Maximum Likelihood and Bayesian Model Fitting

To understand how a drug interacts within the body, we can either simulate populations of individuals and dosage regimens or utilize existing patient data to estimate our parameters. Transitioning from a simulation approach to a model estimation approach is simply the change of the verb applied to the model object. The follow code demonstrates how moving between simulation and fitting in Pumas is a natural transition:

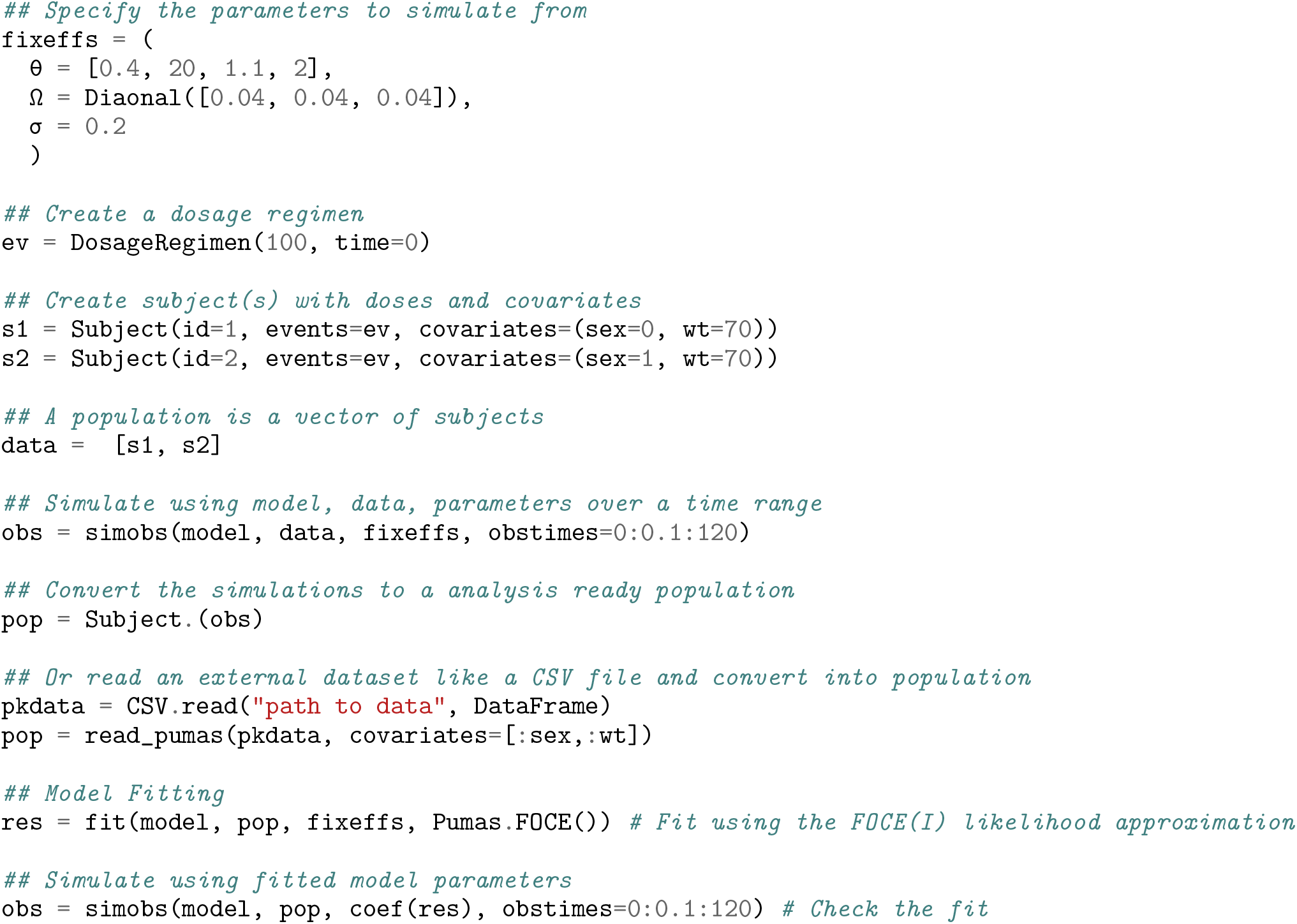

This example first generates a data set using simobs and then proceeds to fit the parameters of the model to data using fit. For fitting one currently has a choice between using:

1. FO: a first order approximation to the likelihood integral
2. FOCE: a first order conditional estimation of the likelihood integral with automatic detection of whether to use interaction or not (FOCEI)
3. LaplaceI: a second order Laplace approximation of the likelihood integral with interaction
4. BayesMCMC: A Bayesian Markov Chain Monte Carlo estimation of model posteriors using the Hamiltonian Monte Carlo no-U-turn sampler (NUTS)^1^
5. SAEM: Stochastic Approximation of the Expectation Maximization algorithm. Requires a slightly different model specification @emmodel.

Note that @emmodel models are required for SAEM and can be used to accelerator other estimation methods. We use the Optim.jl package [27] for the numerical optimization and default to using a safeguarded BFGS algorithm with a backtracking line search for the fixed effects and Newton’s method with a trust region strategy for the Empirical Bayes estimates. The result of the fitting process is a `FittedPumasModel`which we call res. This object can be inspected to determine many quantities such as confidence intervals (as demonstrated later). Importantly, `coef(res)` returns the coeficients for the fixed effects in a `NamedTuple` that directly matches the style of `fixeffs`, and thus we demonstrate calling `simobs` on the returned coeficients as a way to check the results of the fitting process. When applied to the model of Section 1.1, this process returns almost exactly the coeficients used to generate the data, and thus this integration between the simulation and fitting models is routinely used within Pumas to generate unit tests of various methodologies.

## 2 Generalized Quantitative Pharmacology Models with Pumas

The following sections detail a few important features of the Pumas workflow. In summary:

1. We show how the integrated Non-compartmental Analysis (NCA) module can be mixed with NLME simulation and estimation.
2. We show how alternative dynamical formulations, such as stochastic differential equation can be directly utilized in the NLME models.
3. We showcase how the Pumas generalized error distribution form allows for mixing discrete and continuous data and likelihoods
4. We detail the model diagnostics and validation tools
5. We demonstrate the functionality for optimal design of experiments.

### 2.1 Integrated Non-compartmental Analysis (NCA)

NCA is a set of common analysis methods performed in all stages of the drug development program, preclinical to clinical, that has strict rules and guidelines for properly calculating diagnostic variables from experimental measurements [11]. In order to better predict the vital characteristics, Pumas includes a fully-featured NCA suite which is directly accessible from the nonlinear mixed effects modeling suite. Appendix Figures 13 and 14 demonstrate that Pumas matches industry standard software such as PKNCA [9] and Phoenix on a set of 13104 scenarios and 78,624 subjects as described in Section 5. The following code showcases the definition of a NLME model where the observables are derived quantities calculated through the NCA suite, demonstrating the ease at which a validated NCA suite can be coupled with the simulation and estimation routines.

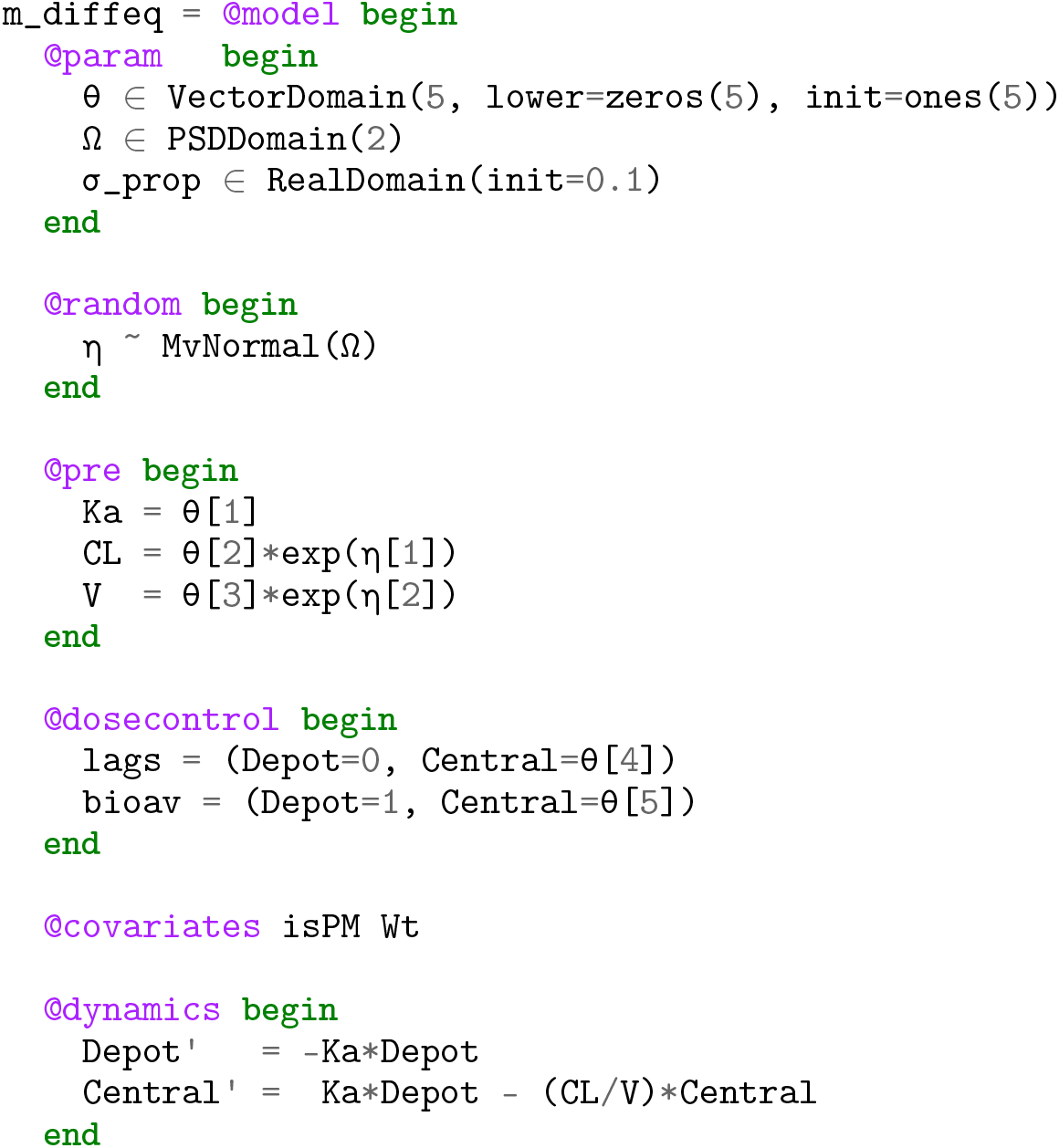

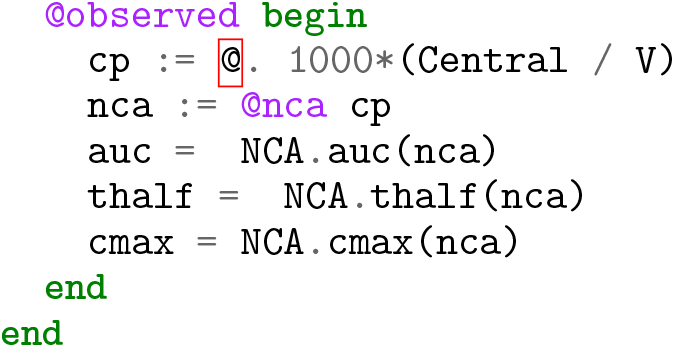

**Figure 13:**
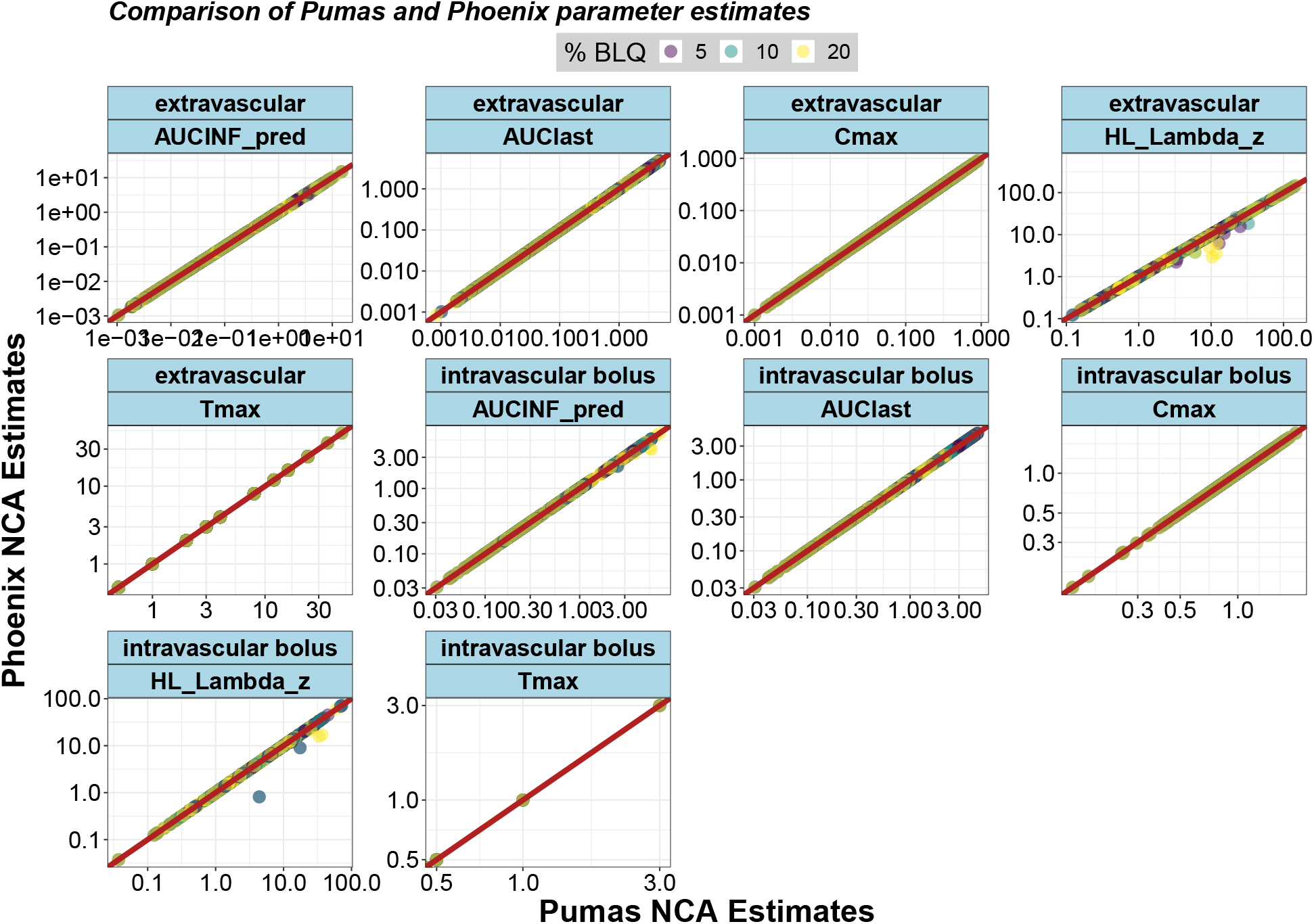
Pumas and Phoenix NCA Results Comparison. Eighteen percent of subjects were randomly selected from the 13104 scenarios to form a subset of 2367 subjects. This smaller subset was used to perform the NCA calculations across the three software while ensuring to maintain the same default options. All analysis were conducted on the same laptop.

**Figure 14:**
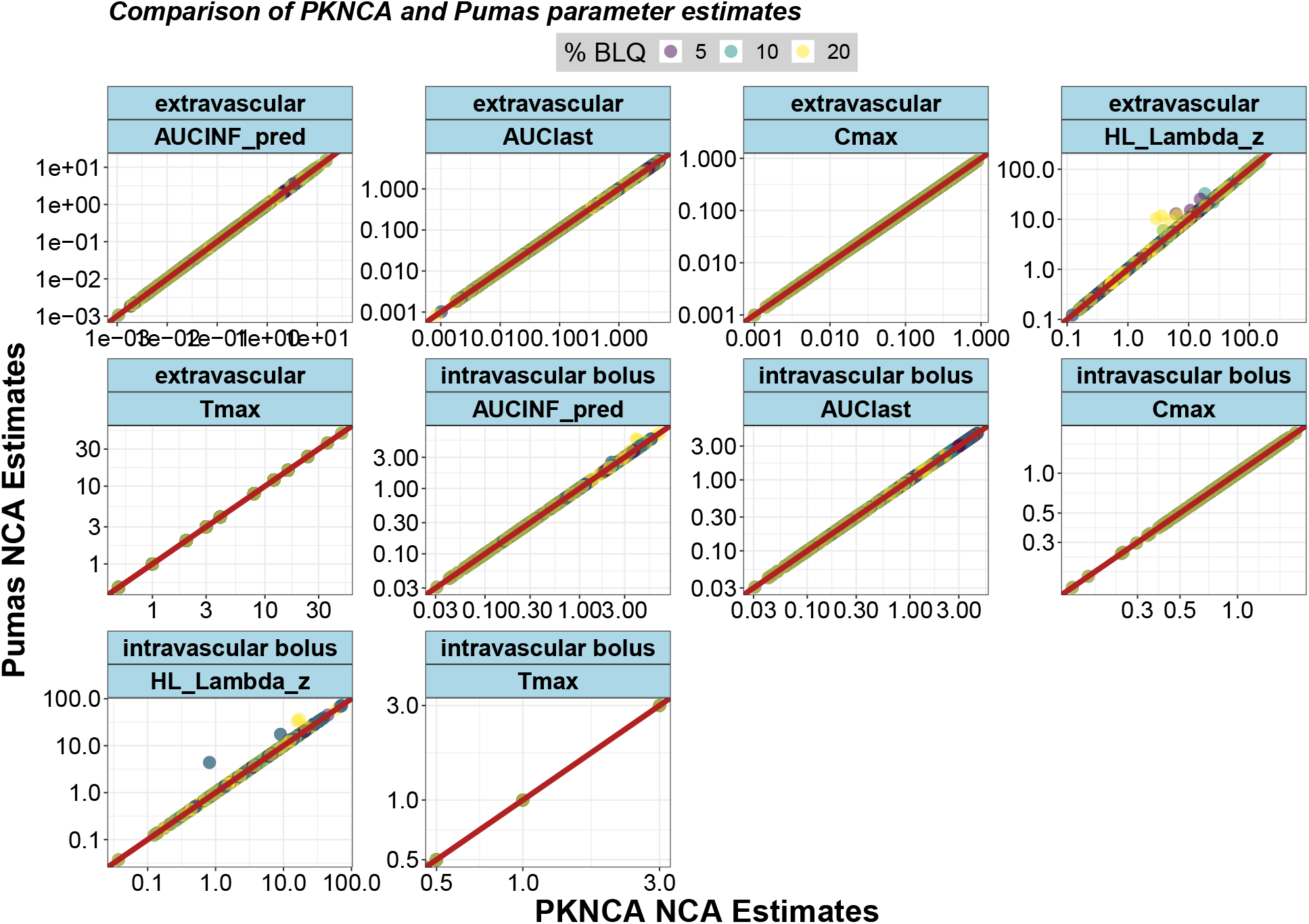
Pumas and PKNCA Results Comparison. Eighteen percent of subjects were randomly selected from the 13104 scenarios to form a subset of 2367 subjects. This smaller subset was used to perform the NCA calculations across the three software while ensuring to maintain the same default options. All analysis were conducted on the same laptop.

### 2.2 Alternative differential Equations via A Function-Based Interface: Stochastic, Delay, and differential-Algebraic Equations for Pharmacology

While ODEs are a common form of dynamical model, in many circumstances additional features are required in order for the realism of the model to adequately predict the behavior of the process. For example, many biological processes are inherently stochastic, and thus an extension to ordinary differential equations, known as stochastic differential equations, takes into account the continuous randomness of biochemical processes [34, 37, 22, 49]. Many biological effects are delayed, in which case a delay differential equation which allows for an effect to be determined by a past value of the system may be an appropriate model [19, 23]. Together, Pumas allows for the definition of dynamical systems of the following non-standard forms:

1. Split and Partitioned ODEs (Symplectic integrators, IMEX Methods)
2. Stochastic ordinary differential equations (SODEs or SDEs)
3. differential algebraic equations (DAEs)
4. Delay differential equations (DDEs)
5. Mixed discrete and continuous equations (Hybrid Equations, Jump Diffusions)

where each can be simulated with high-performance adaptive integrators with specific method choices stabilized for stiff equations. The following code demonstrates the definition solving of a stochastic differential equation model with steady state dosing via a high strong order adaptive SDE solver specified using an alternative interface known as the Pumas function-based interface. This interface is entirely defined with standard Julia functions, meaning that any tools accessible from Julia can be utilized in the pharmacological models through this context.

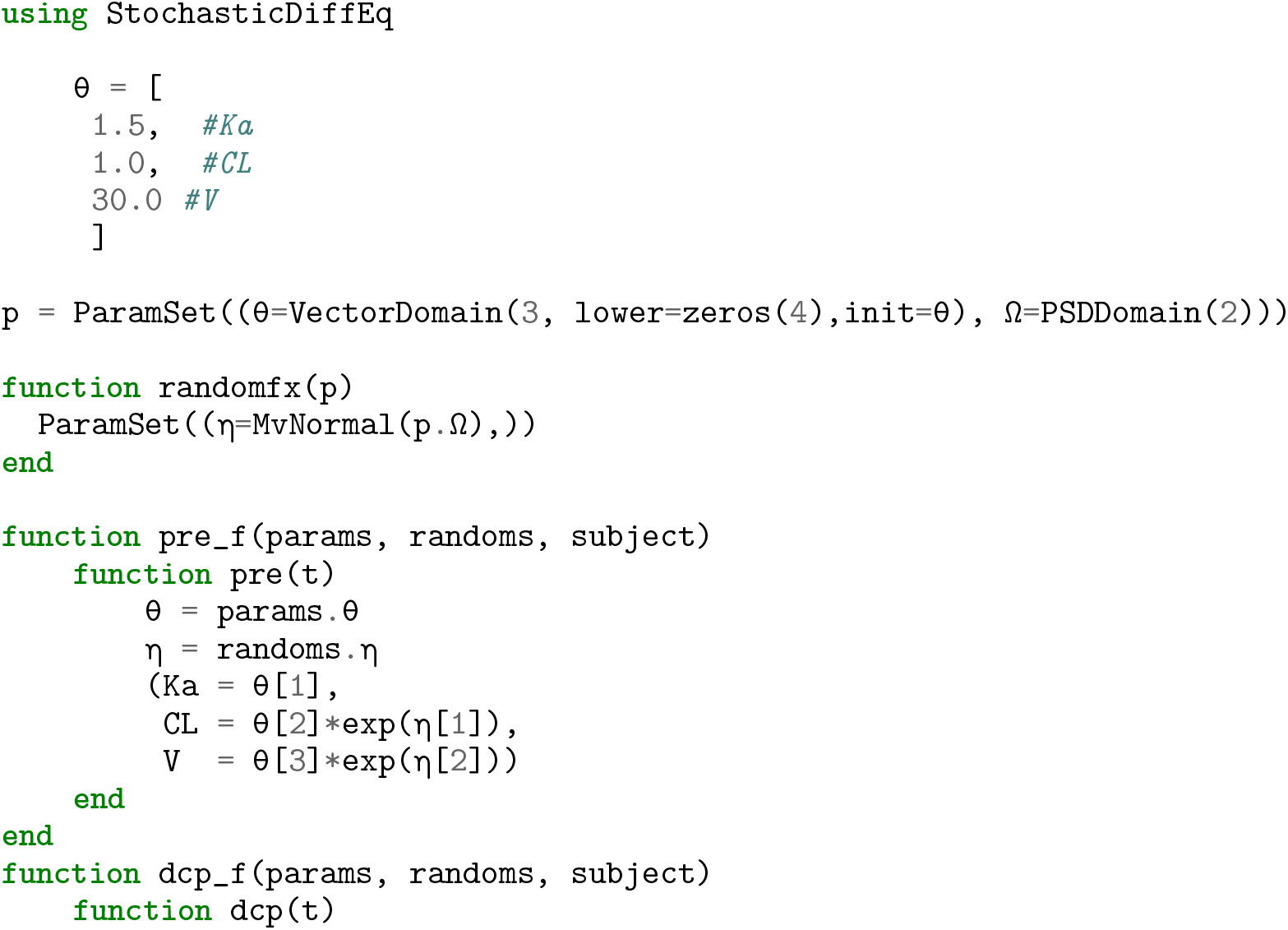

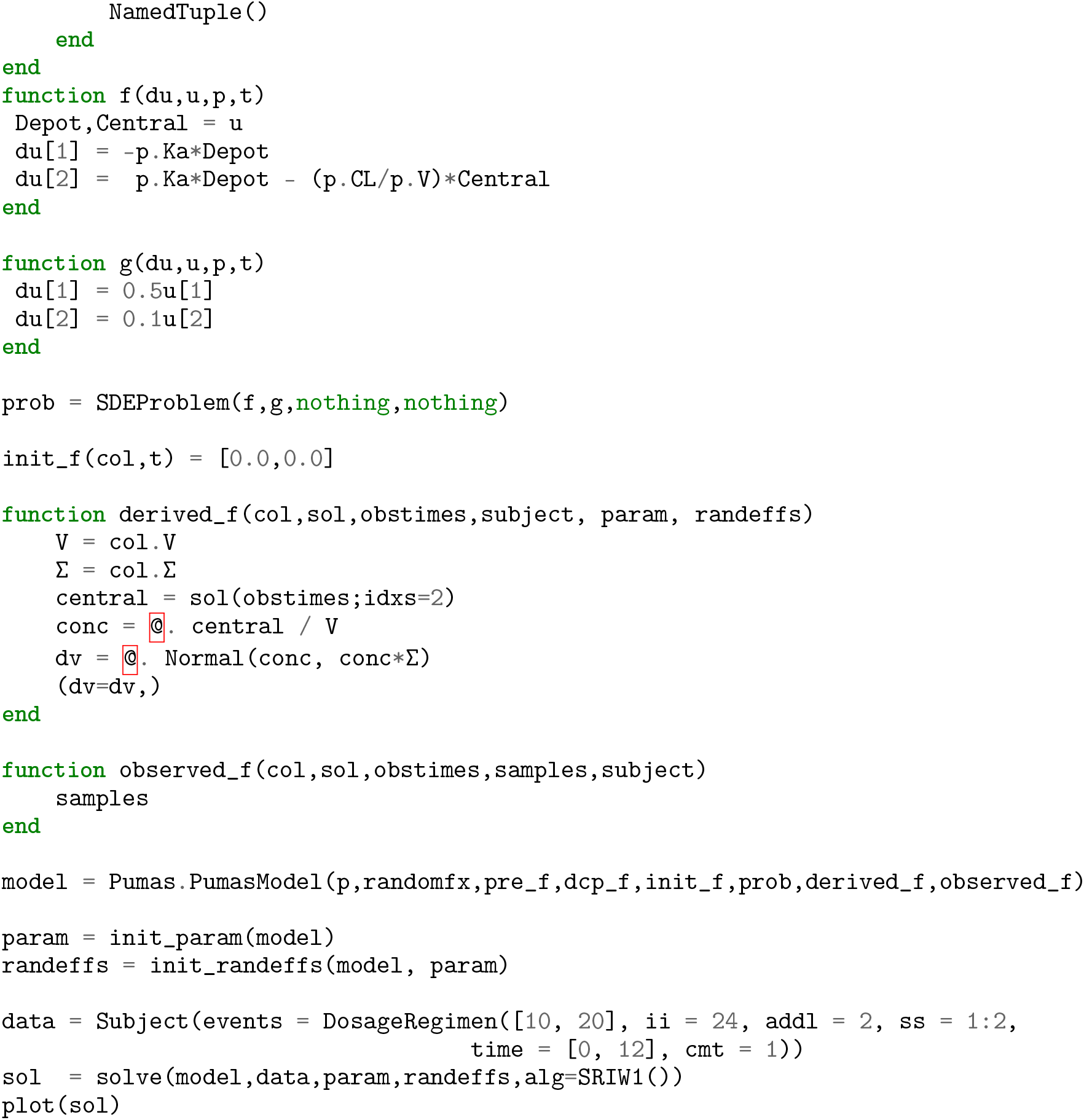

### 2.3 Generalized Error Distribution Abstractions

Traditional nonlinear mixed effects models formulate the error distribution handling in a different manner, focusing on the perturbation via the error against the estimate. Mathematically, this amounts to having an *ϵ* measurement noise, i.e. defining the observables as:

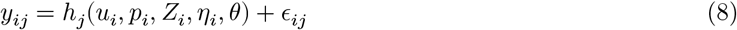

Instead of using the perturbation format, Pumas allows for users to specify the likelihood directly via any arbitrary Distributions.jl [5] distribution, with the mean of the distribution corresponding to the prediction. This has some immediate advantages. For one, many distributions are not directly amenable to the perturbation form. For example, one may wish to model a discrete observable, such as a pain score, probabilistically based on clinical outputs. A discrete likelihood function, like a Poisson distribution, can thus be given to described the predicted outputs like in the following example:

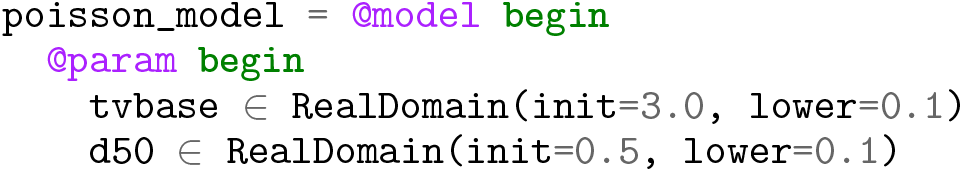

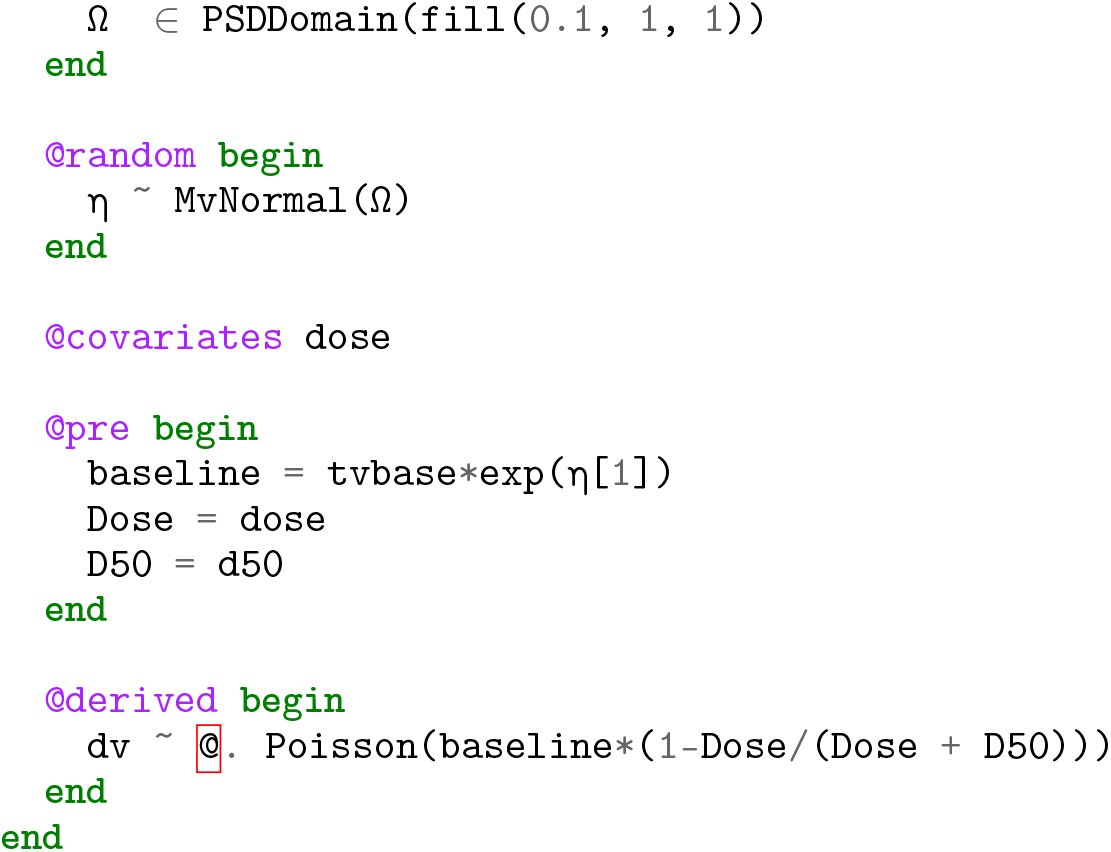

This type of model could not be described with continuous perturbations and thus would have to be approximated in other scenarios. In addition, since any distribution is possible in this format, users can extend the modeling schema to incorporate custom distributions. Discrete Markov Chain models, or Time to Event models for example can be implemented as a choice of a custom distribution, thus making it easy to extend the modeling space directly from standard language use.

### 2.4 Integrated and Parallelized Model Diagnostics and Validation

After running the estimation procedure, it is important to have a wide suite of methods to post-process the results and validate the predictions. Model diagnostics are used to check if the model fits the data well. Pumas provides a comprehensive set of diagnostics tooling A number of residual diagnostics are available as well as shrinkage estimators. Additionally, for model validation, Visual Predictive Checks (VPCs) [18] can be used for a variety of models in a performant and robust manner.

The diagnostics tooling comprises of:

1. Residuals: Populations residuals take the inter-individual variability into account as opposed to Individual residuals which are correlated due to the lack of accounting for inter-individual variability. The following Population residuals are available in Pumas^2^:

- Normalized Prediction Distribution Errors (NPDE) [26]
- Weighted Residuals (WRES)
- Conditional Weighted Residuals (CWRES)
- Conditional Weighted Residuals with Interaction (CWRESI)

Similarly the following Individual residuals are also available -

- Individual Weighted Residuals (IWRES)
- Individual Conditional Weighted Residuals (ICWRES)
- Individual Conditional Weighted Residuals with Interaction (ICWRESI)
- Expected Simulation based Individual Weighted Residuals (EIWRES)
2. Population Predictions: Population prediction are defined as the difference between the observations and the model expectation for subject *i*, i.e. *y*_*i*_ – **E**[*y*_*i*_] where the **E**[*y*_*i*_] are the population predictions. The population predictions, EPRED, PRED, CPRED and CPREDI are defined as the **E**[*y*_*i*_] from equations of NPDE, WRES, CWRES and CWRESI respectively.
3. Individual Predictions: The individual residuals are defined as the difference between the observations and the model prediction for subject *i*, i.e. 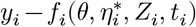 where the 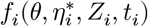 are the individual predictions. The individual predictions IPRED, CIPRED and CIPREDI are defined as the 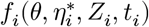 from equations of IWRES, ICWRES and ICWRESI. The individual expected prediction (EIPRED) is defined as the individual mean conditioned on *η*_*k*_, i.e. **E**[*y*_*i*_|*η*_*k*_], from equation of EIWRES.
4. Akaike Information Criterion (AIC) and Bayesian Information Criterion (BIC)
5. *η* Shrinkage and *ϵ* Shrinkage [36]

The above discussed diagnostics can be evaluated through the `inspect`function. Additionally, the `infer` function is available for computing the covariance matrix of the population parameters either using the sandwich estimator or the inverse hessian approximation thus giving the the 95% confidence intervals and the standard errors for the parameter estimates. The inferfunction can also be used to perform `Bootstrapor SIR`(sampling importance resampling). As an example below, we call `infer` and `inspecton` the `FittedPumasModel` object `res` obtained as the result of `fit` call in earlier section.

~~~
resinfer = infer(res)
DataFrame(resinfer)
resinpect = inspect(res)
DataFrame(resinspect)
~~~

Additionally several out of the box visualizations are available for practitioners to evaluate the model fit that are mainly provided by the PumasUtilities.jl package. Convergence plot for fitting obtained with `convergence`.In Pumas we provide the ability to run Visual Predictive Checks with `vpc` function. For continuous models VPCs are computed using the Quantile Regression based approach discussed in [20]. The syntax for computing and plotting the VPCs is shown below followed by the Figure 3 of a stratified VPC plot in Pumas.

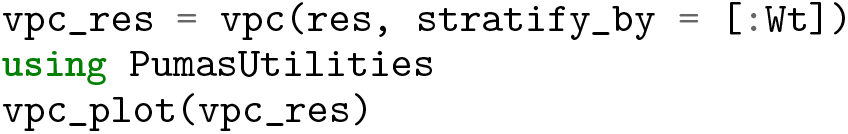

### 2.5 Optimal Design of Experiments

Optimal design of clinical trials has increasingly played a central role in increasing the effectiveness of such studies while minimizing costs [25]. To facilitate the advancement of optimal design methodologies into standard clinical trial workflows, Pumas includes functionality for high performance calculations of the Fisher Information Matrix (FIM) which is central to optimal design methodologies. The function `FIM` takes in a `PumasModel`, a `Subject`and a set of model parameters as input and returns the expected FIM using a normality approximation [43]. The following demonstrates the workflow for calculating the expected FIM in the Warfarin PK model [29] at specific sample times t:

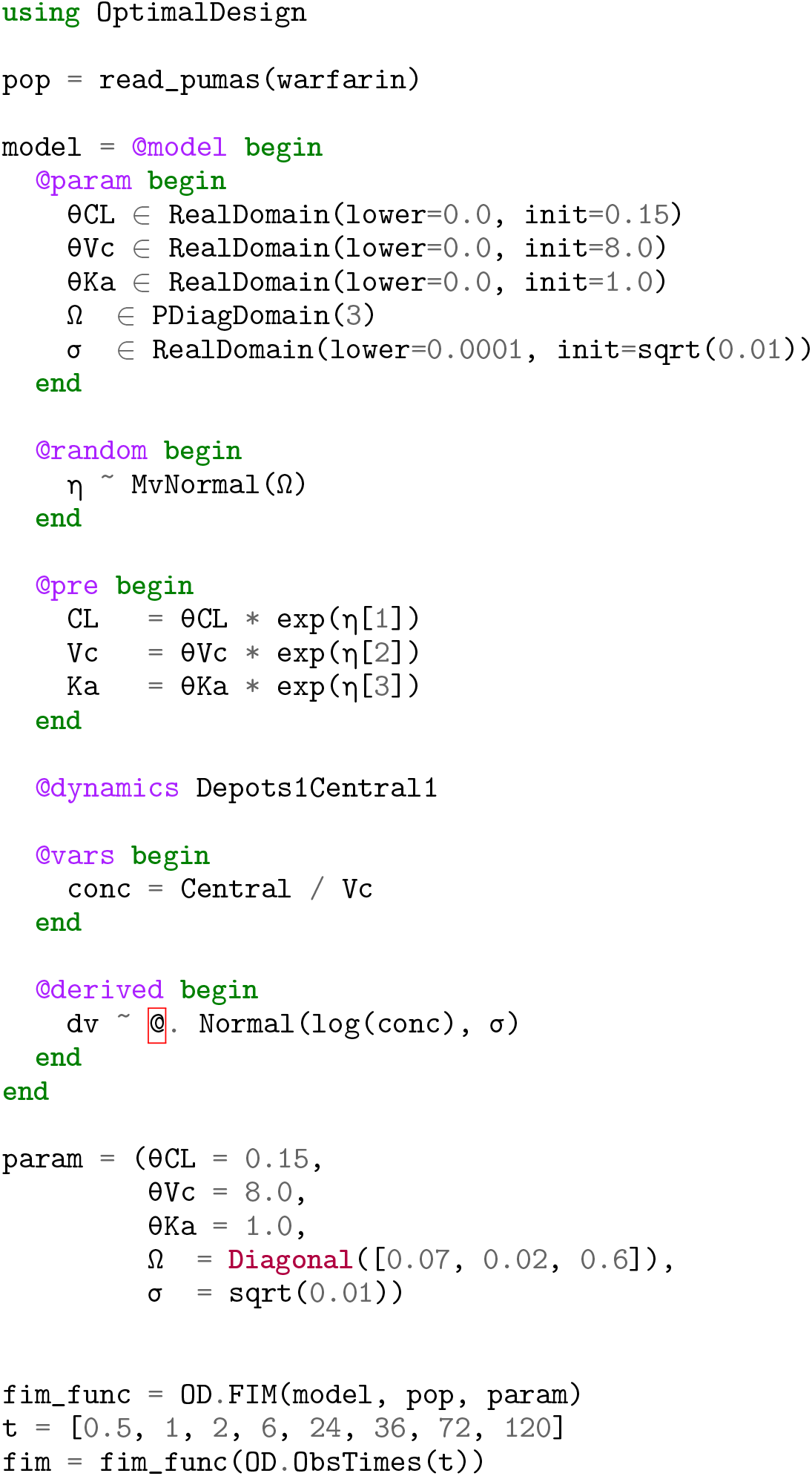

Note that the derivative calculations used within the FIM calculation are the discrete sensitivity analysis derivatives derived via forward-mode automatic differentiation as described in Section 3.2. Finite difference approximation of derivatives is also optionally available. The following script further demonstrates how to do sample time optimization using 4 additional lines of code given a `model`, a subject or population `data`, and fixed effects `param`:

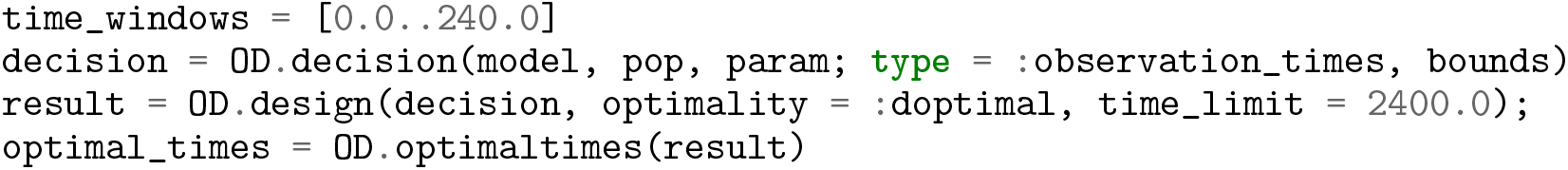

Note that the same Pumas subjects and models can be used to perform optimal design thus giving a streamlined user experience. For more information on the various optimal design tasks that can be performed and options, please refer to the Pumas documentation.

### 2.6 Post-Processing Utilities

A wide selection of post-processing utilities are provided together with Pumas that integrate with the rest of the software system to provide users with essential insights into their data, models, and results. These take the form of a comprehensive set of plotting functions for visualizing user data at each stage of an analysis including goodness-of-fit, VPC, and covariates plots. Figure 4 showcases one of the aforementioned goodness-of-fit plots. In addition, several interactive web-based applications for exploring and evaluating your results are also provided, as well as automated static reporting to generate standardized reports containing tables, listings, and figures based on a user’s results. Figure 5 showcases the initial parameter estimates web application for visual tuning of parameter estimates. Pumas has an ever increasing set of such post-processing tools, and thus consult the current documentation for a complete list.

## 3 High Performance and Stability-Enhanced Model Fitting

### 3.1 Fine-Tuning differential Equation Solver Behavior to Model Features

The core computation of the model fitting process utilizes the `simobs` function for generating solutions to the differential equation during the likelihood approximation, meaning that every step of the optimization is solving thousands of the same small differential equation representing different possible parameter configurations amongst all subjects in a clinical trial. Thus, while in isolation these small ODEs may simulate very fast, real-world NLME model fitting with large numbers of subjects consistently arrives at workflows which take hours to days or weeks with the majority of the time due to the cost of solving small ODEs. Thus practical workflows of industry pharmacologists would be heavily impacted if the speed of these systems could be dramatically decreased.

Pumas recognizes the crux of the computational issue and thus has many new features for optimizing the internal solve of the fitting process. The options for controlling the solvers are same between the simulation and estimation workflows. The full gamut of options from differentialEquations.jl are exposed to allow users to control the solvers as much as possible. This allows for specializing the solver behavior on the known characteristics of the functions and its solution. For example, the concentrations modeled in the ODEs need to stay positive in order for the model to be stable, but numerical solvers of ODEs do not generally enforce this behavior which can cause divergences in the optimization process. In Pumas, one can make use of advanced strategies [39] like rejecting steps out of the domain by using `isoutofdomain` or using the `PositiveDomain` callback.

However, a more immediate effect of this connection is the ability to choose between a large set of highly optimized integration methods. Table 1 shows timing results of Pumas on pharmacokinetic (PK) and pharmacokinetic/pharmacodynamic (PK/PD) models using both the native differentialEquations.jl methods and some classic C++ and Fortran libraries and demonstrates a performance advantage around 2x-440x (mean of 102x with median 3.7x) between the best Julia-based method against the best wrapped C++ or Fortran solver method. This table also demonstrates a few different dimensions by which this performance advantage is achieved. First of all, the differentialEquations.jl library uses different Runge-Kutta methods which are derived to have asymptotically better error qualities for the same amount of work [45, 47], with a much larger effect for the high order (7th order) integrator. Secondly, differentialEquations.jl uses a tuned PI-adaptive timestepping method [42] which is able to stabilize the solver and increase the step sizes, thus decreasing the total amount of work to integrate the equation. Lastly, Pumas is able to utilize the JIT compiler to compile a form of the differential equation solver that utilizes stack-allocated arrays and is specific to the size and ODE function. While this optimization only applies to small ODE systems (the optimization is no longer beneficial at around 10 or more ODEs), many pharmacometric models fall into this range of problems and these benchmarks demonstrate that it can have a noticeable effect on the solver time.

**Table 1:** effect of specialized ODE solvers on forward simulation of small pharmacometric ODE models. Shown is the effect of the ODE solver choice on the speed of the forward pass of common pharmacometric models. OC for the one-compartment model of Section 1.1, MR for the multiple response model described in Section 5, and HCV for the hapatitis C model described in Section 5. Times are all shown in seconds. By default the tolerances for the solvers was 1 × 10^−^3 for the relative tolerance and 1 × 10^−^6 for the absolute tolerance, while LT stands for low tolerance with 1 × 10^−^8 for the relative tolerance and 1 × 10^−^12 for the absolute tolerance (representative of tolerances used when simulating vs when fitting parameters). Solutions were checked against a tolerance 1 × 10^−14^ reference solution to ensure the actual errors were within the same order of magnitude. The automatic stiffness detection AutoTsit5-Rosenbrock23 method is a combination between Tsit5 [45] and a Rosenbrock method Rosenbrock23 [38], while Tsit5 and Vern7 [47] are explicit Runge-Kutta methods from the differentialEquations.jl library [33]. CVODE is from the Sundials library [17], while DOP853 and DOPRI are from the Hairer Fortran methods suite [13]. The Array methods are generically compiled for heap-allocated mutable arrays, where the SArray versions are specially optimized for the specific size of the ODE using Julia’s JIT compiler.

| ODE Solver vs Time (seconds) | OC | OC(LT) | MR | MR(LT) | HCV | HCV(LT) |
| --- | --- | --- | --- | --- | --- | --- |
| AutoTsit5-Rosenbrock23 (SArray) | $9.45318 \times 10^{-7}$ | **$9.39381 \times 10^{-7}$** | $7.8115 \times 10^{-5}$ | 0.00040444 | 0.000119592 | 0.000610029 |
| Tsit5 (SArray) | **$9.36826 \times 10^{-7}$** | $9.46667 \times 10^{-7}$ | **$7.2851 \times 10^{-5}$** | 0.00038857 | **0.000108575** | 0.000546279 |
| Vern7 (SArray) | $9.43423 \times 10^{-7}$ | $9.4036 \times 10^{-7}$ | 0.000103155 | **0.000289059** | 0.000156718 | **0.000476379** |
| AutoTsit5-Rosenbrock23 (Array) | $7.9365 \times 10^{-5}$ | 0.00035122 | 0.000100629 | 0.000391078 | 0.000146214 | 0.000671795 |
| Tsit5 (Array) | $6.5564 \times 10^{-5}$ | 0.000322259 | $7.6432 \times 10^{-5}$ | 0.000366739 | 0.000127374 | 0.000637998 |
| Vern7 (Array) | $9.6297 \times 10^{-5}$ | 0.000249384 | 0.00012611 | 0.000292689 | 0.000196665 | 0.000559559 |
| CVODE (Array) | 0.000149114 | 0.000411655 | 0.000168402 | 0.00191018 | 0.000546243 | 0.00144688 |
| DOP853 (Array) | 0.000223506 | 0.000968525 | 0.00023595 | 0.00105208 | 0.000468118 | 0.00242868 |
| DOPRI (Array) | 0.000255584 | 0.000467723 | 0.000293277 | 0.000575954 | 0.000580232 | 0.00183609 |

One additional advantage of this tweak-ability is the ease to span multiple domains. Physiologically-based pharmacokinetic (PBPK) models are typically larger stiff ODE-based models which incorporates systems-type mechanistic modeling ideas to enhance the model’s predictive power [24]. Table 2 demonstrates the performance advantage of the native Julia methods that are unique to Pumas on a 15 stiff ODE PBPK model with steady state dosing, demonstrating a 2x-4x performance advantage over the classic CVODE method used in many other pharmacometrics modeling suites. We note that the benchmark platform gave pessimistic estimates for the speedup of the Julia-specific tools, with some computers showing a 4x-5x advantage over the classic methods as demonstrated in the Appendix.

**Table 2:** effect of specialized ODE solvers on forward simulation the Voriconazole physiologically-based pharmacokinetic (PBPK) model [51]. Shown is the effect of the ODE solver choice on the speed of the forward pass of the PBPK model with steady state dosing. Given the high degree of variance in the stiff ODE solver accuracies at the same tolerances, the tolerances were aligned using a reference solution computed at 1 × 10^−14^ tolerances, and high tolerance was chosen to be the tolerance pair by power of 10 which achieve a true error closest to 1 × 10^−5^ and for low tolerance the pair closes to 1 × 10^−9^ The automatic stiffness detection method AutoVern7-Rodas5 is a combination between Vern7 and Rodas5, while Rodas5 [48], KenCarp4 [21], and RadauIIA5 [14] are implicit methods from the differentialEquations.jl library [33]. CVODE is from the Sundials library [17]

| ODE Solver | PBPK | PBPK(LT) |
| --- | --- | --- |
| AutoVern7-Rodas5 | 0.147 | **0.520** |
| Rodas5 | **0.141** | 1.003 |
| KenCarp4 | 0.250 | 3.224 |
| RadauIIA5 | 0.369 | 3.467 |
| CVODE | 0.519 | 1.029 |

### 3.2 Fast and Accurate Likelihood Hessian Calculations via Automatic differentiation

When performing maximum likelihood estimation or Bayesian estimation with a gradient-based sampler like Hamiltonian Monte Carlo, the limiting step is often the calculation of the gradient of the likelihood. Finite difference calculations are not efficient since every calculation of a perturbation involves a numerical solve of the ODE system and either or two perturbations are required for each model parameter for forward and central differencing respectively. Furthermore, the finite difference approximation of a derivative is an unstable process [10] which in some cases can result in very inaccurate gradients when combined numerical solutions to ODEs.

The performance of the derivative calculations for the marginal likelihoods can be improved by utilizing a formulation with the sensitivity equations due to [1]. While very efficient, the method relies on second order derivatives in a way that makes the process unstable and accurate derivatives are required for process to work well. Instead of utilizing the traditional form of the sensitivity equations, Pumas generates an implementation of discrete forward sensitivity analysis by utilizing dual number arithmetic through the differential equation solver [35]. Our group has previously shown that this discrete sensitivity analysis via automatic differentiation outperforms traditional sensitivity analysis since it allows the compiler more freedom to optimize the generated code for passes like single instruction, multiple data (SIMD) auto-vectorization [32].

Table 3.2 demonstrates the effect on run time and and accuracy of using the sensitivity method from [1] with finite difference and automatic differentiation based derivatives respectively as well as a simple but expensive finite difference based gradient computation. The error is computed relative to a solution computed with 256 bit precision floating point numbers. All ODE solutions are computed to a relative tolerance of 10^−8^. The results show that the simple finite difference based gradient doesn’t have too large an error but is slow while the faster sensitivity based method has a large relative error. The error is so large that gradient based optimization is no longer practical. In contrast, the sensitivity based gradient computation based on automatic differentiation loses almost no precision and is even faster that the finite difference based gradient because it requires fewer evaluations of the objective function.

**Table 3:** Timings and relative errors for the gradient computations of the marginal likelihood in a non-linear fixed effects model with time varying coefficients in the ODE. Shown are the timings of the three likelihood gradient calculation methods along with their relative error. AD stands for automatic differentiation while FD stands for finite differentiation, demonstrating both the efficiency and accuracy gains obtained through by utilizing the automatic differentiation derived discrete sensitivity analysis.

|  | AD Almquist | FD Almquist | FD simple |
| --- | --- | --- | --- |
| Relative error | **9.52e-7** | 0.811 | 0.00047 |
| Time in seconds | **0.431** | 0.641 | 18.6 |

**Table 4:** The table shows the time comparison when evaluating the determinant of the expected Fischer information matrix in both Pumas and PopED [30].

| Model | Pumas | PopED |
| --- | --- | --- |
| Warfarin | **0.187** ms | 1.834 ms |
| HCV | **480.89** ms | 20279.22 ms |

### 3.3 Accelerated Maximum Likelihood Fitting

Maximum likelihood directly corresponds to repeated calculation of likelihood gradients. Given the performance advantages that are obtained due to the ODE solver and discrete sensitivity analysis advantages over previous software, one would predict that the model estimation routines would see a similar benefit as derived from these components. Appendix 5 describes the full set of benchmark models for maximum likelihood estimation. In Figure 6 we demonstrate this is the case by calculating the elapsed run time to estimate 6 models in both Pumas and NONMEM with FOCE/LaplaceI and SAEM (an EM based approach). We see a 2x-140x performance advantage for Pumas (mean 78x, median 81x), growing as the complexity of the problem increases, similar to the difference between the non-fully optimized ODE solver and the alternative array-based algorithms and implementations. We note that Pumas has the ability to automatically detect analytical solutions of ODEs to further accelerate the solving of these models, though this feature was turned off to ensure fairness in the benchmarking process. Figure 10 benchmarks the same models with multiple dosing, demonstrating similar results as the dosing strategy is changed. Figure 7 demonstrates that this performance advantage over NONMEM extends to larger nonlinear models, with a mean speedup of 80x and median of 82x, showcasing the general applicability of the performance enhancements seen in Pumas. Note that compilation only occurs on the first run and is a property of the code, not runtime values (such as number of subjects or dosing strategy), and thus has its cost as relatively constant as the runtime cost of a model grows. Thus, this figure also showcases that the aforementioned JIT compilation strategy leads to substantial improvements as the cost of the fitting problem increases. Appendix 5 validates that the Pumas and NONMEM estimations arrive at the same values, demonstrating that acceleration is achieved without loss of accuracy.

### 3.4 Automated Parallelism and Full-Application Benchmarks

Pumas is able to automatically parallelize the solution of the NLME model solution across subjects. 2 forms of parallelism are currently available:

1. Multithreaded parallelism for shared memory machines
2. Multiprocessing for distributed memory architectures

Multithreaded is enabled by default and no extra steps are required for this parallelism to occur. Multi-processing is possible using the distributed computing functionality in Pumas. Figure 8 demonstrates the good scaling of multithreading on a PK/PD maximum likelihood estimation. Since Pumas is built on the JuliaHub cloud computing platform it is simple to scale the available computational power up and down to essentially arbitrary numbers of cores. The user simply chooses how many cores and memory they want to have available when starting their instance.

### 3.5 High Performance Non-Compartmental Analysis (NCA)

In the previous sections we demonstrated that the integrated NCA suite reproduced the results of industry-standard tools, demonstrating the correctness of the implementation over 13104 scenarios. In addition to determining the correctness, we calculated the run times. The full analysis tool 3 hours and 16 minutes in PKNCA while in Pumas the run time was 56 seconds, demonstrating a 210x acceleration. This speed is particularly useful when 1,000s of clinical trials are simulated during the planning of a large patient trial.

### 3.6 Accelerated Optimal Design of Experiments

In order to assess the effectiveness of accelerated differential equation solving on optimal design workflows, we tested the calculation time of the FIM on the Warfarin PK model (Section 2.5) and the HCV PK/PD model Section 5 against the PopED Population Optimal Experimental Design framework [30]. Table 3.6 showcases that Pumas calculates the FIM approximately 10x faster than PopED on the Warfarin model and approximately 42x faster on the HCV model. Much of the acceleration can be attributed to the `AutoVern7(Rodas5())` ODE solver used in the Pumas version of the FIM calculation as opposed to the ode45 method internally used by PopED. This demonstrates how central improvements to the solving of small ODEs can give large advantages to real applications built on top of such methodologies.

**Table 5:** Voriconazole PBPK results on AMD Ryzen 9 5950X 16-Core Processor.

| ODE Solver | PBPK | PBPK(LT) |
| --- | --- | --- |
| AutoVern7-Rodas5 | 0.0561175 | **0.197751** |
| Rodas5 | **0.055821** | 0.384349 |
| KenCarp4 | 0.329876 | 1.13387 |
| RadauIIA5 | 0.1496 | 1.34265 |
| CVODE | 0.285117 | 0.818587 |

**Table 6:** Voriconazole PBPK results on 28 × Intel(R) Core(TM) i9-9940X CPU @ 3.30GHz.

| ODE Solver | PBPK | PBPK(LT) |
| --- | --- | --- |
| AutoVern7-Rodas5 | 0.0773158 | **0.261663** |
| Rodas5 | **0.0732951** | 0.518652 |
| KenCarp4 | 0.136943 | 1.66849 |
| RadauIIA5 | 0.150421 | 2.40308 |
| CVODE | 0.247077 | 0.522646 |

**Table 7:** Voriconazole PBPK results on 64 × AMD EPYC 7513 32-Core Processor.

| ODE Solver | PBPK | PBPK(LT) |
| --- | --- | --- |
| AutoVern7-Rodas5 | 0.073701 | **0.242261** |
| Rodas5 | **0.0697943** | 0.50318 |
| KenCarp4 | 0.134364 | 1.6422 |
| RadauIIA5 | 0.175766 | 0.716251 |
| CVODE | 0.254783 | 0.538172 |

Beside single FIM evaluation, we also ran a sample time optimization optimal design problem using the IVGTT insulin model [40, 41]. The problem involved the optimization of 10 unique sample times over a time window from time 0 to 240. Each sample was made of 3 sub-samples except the first one which only had 2 sub-samples. There was 1 unique subject in the study repeated 42 times and all the subjects were forced to have identical sample times. The log determinant of the expected FIM was maximized in the optimization. The same differential equation tolerance of 10^−5^ was used in both software. The optimal design problem was ran using Pumas and PopED for 1 hour starting from the same initial design and the profile of the objective function profiles were plotted with respect to time as shown in figure 9.

## 4 Conclusion

Pumas pulls together a diverse array of pharmacometrics problems into a single platform and utilizes the JIT optimization to directly specialize the internal solvers. We have showcased how the Pumas platform achieves across the board performance improvements from early to late stage clinical analyses over existing pharmacometrics tools. This more broadly illustrates how integrating optimizing compilers into dynamic tool chains can improve performance over traditional approaches which do not specialize on the problem’s scale. Even something as standardized as an ODE solver can be improved. Pumas is already in production use even at this stage of early development and will continue to increase its performance as it demonstrates new features to the pharmacometrics community. Indeed, the results on the large nonlinear dynamics estimation problems in comparison to the ODE solver results suggests more potential performance improvements can be had by tuning the optimization schemes. Together, the innovations of this new industry-ready software will facilitate unprecedented acceleration and scalability in pharmaceutical modeling and simulation leading to increased efficiency in drug development and precision in real-time personalized healthcare delivery.

## 5 Acknowledgements

The Pumas project was started in 2017 with the support of the University of Maryland Baltimore School of Pharmacy (UMB). Professor Jill Morgan, the chair of Pharmacy Practice and Science (PPS) and Dean Natalie Eddington were instrumental in their support. The staff, faculty, students and researchers at the Center for Translational Medicine in UMB drove a lot of the initial testing and we are extremely thankful for their patience while adopting a new language and tool. Brian Corrigan from Pfizer who encouraged us throughout the journey and nudged the team to achieve new heights. All the early adopters and believers. Viral Shah and Deepak Vinchii from Julia Computing for being our trusted technology collaborators. Last but the most important thank you goes out to the members of the Julia Language community who develop world class scientific computing packages and create an environment that is welcoming to newcomers.

## Appendix Benchmark Details

All benchmarks were ran using Pumas 2.1 on an isolated Xeon E5-2698 v3 48 core and 2.3 GHz processor. Comparisons against NONMEM used v7.5. Comparisons against Phoenix used v8.1. Comparisons against PKNCA used v0.9.5. Comparisons against PopED used v2.13.

## PBPK Benchmark on Alternative Computers

These additional benchmarks demonstrate that the shown results follow a common trend, with the demonstrated results being some of the most conservative performance figures for the Julia-based methods.

## Maximum likelihood benchmarks with multiple doses

Shown in Figure 10. The mean speedup was 63x with median 73x.

## Model Definitions

## Multiple Response Model

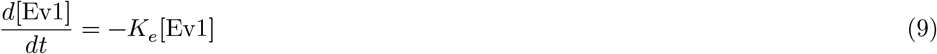

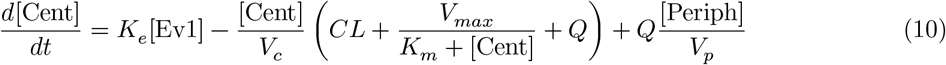

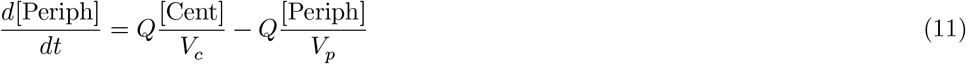

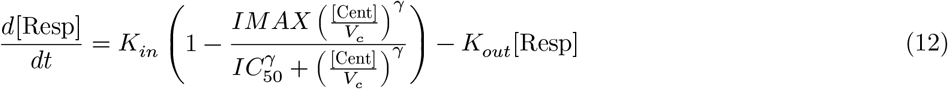

where:

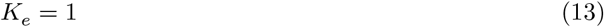

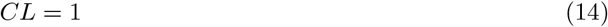

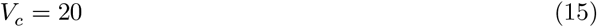

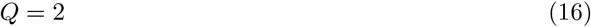

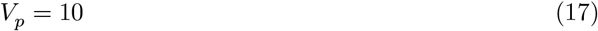

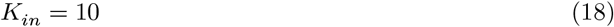

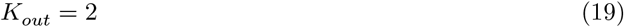

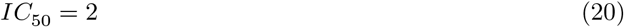

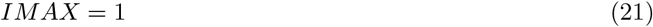

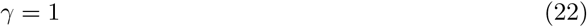

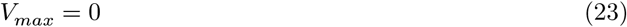

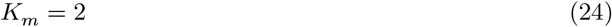

## HCV Model

The “HCV Model” is a multiple response PKPD model from [29] that models the effect of a pegylated interferon dose given as a 24h infusion once a week for 4 weeks. Twelve simulated samples are obtained for the PK and PD at times *t* = [0.0, 0.25, 0.5, 1.0, 2.0, 3.0, 4.0, 7.0, 10.0, 14.0, 21.0, 28.0]. The internal dynamics are:

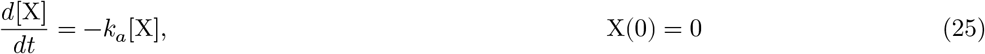

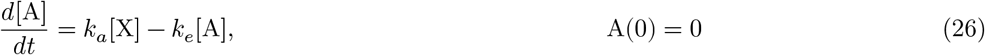

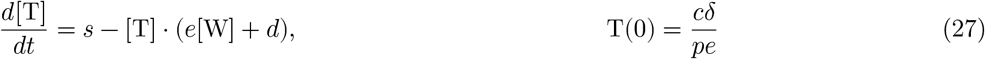

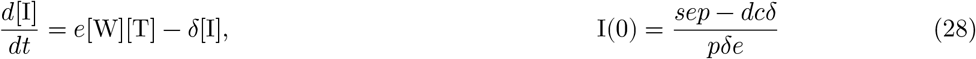

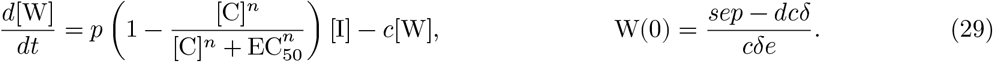

where *C(t)* = *A(t)*/*V*_*d*_. The PK variable *C* is modelled with an additive normal distribution and the PK variable *W* is modelled as being log-normal. We use the parameters reported in the original paper.

## Dosing Regimen Verification

The event handling of Pumas and the accuracy of simulation under different kinds of events was evaluated systematically. The examples for this testing were adapted from mrgsolve test suite^3^. The dosing and sampling scenarios were setup in R’s mrgsolve package which were then used to simulate concentration time profile for a single subject in NONMEM and Pumas. The listing of all the tested scenarios is presented below:

1. Infusion (10 mg/hr) into the central compartment with 4 doses given every 12 hours
2. Infusion (10 mg/hr) into the central compartment with lag time (5 hr) with 4 doses given every 12 hours
3. Infusion (10 mg/hr) into the central compartment with lag time (5 hr) and bioavailability (0.4) with 4 doses given every 12 hours
4. Infusion (10 mg/hr) into the central compartment at steady state (ss)
5. Infusion (10 mg/hr) into the central compartment at steady state (ss) with 81% bioavailability, where frequency of events (ii) is less than the infusion duration (DUR)
6. Infusion (10 mg/hr) into the central compartment at steady state (ss) with 100% bioavailability, where frequency of events (ii) is less than the infusion duration (DUR)
7. Infusion (10 mg/hr) into the central compartment at steady state (ss) with 100% bioavailability, where frequency of events (ii) is a multiple of infusion duration (DUR)
8. Infusion (10 mg/hr) into the central compartment at steady state (ss) with 41% bioavailability, where frequency of events (ii) is exactly equal to the infusion duration (DUR)
9. Infusion (10 mg/hr) into the central compartment at steady state (ss) with 100% bioavailability, where frequency of events (ii) is exactly equal to the infusion duration (DUR)
10. Oral dose at steady state with lower bioavailability of 41%
11. Oral dose at steady state with lower bioavailability of 41% and 5 hour lag time
12. Zero order infusion followed by first order absorption into gut
13. Zero order infusion into central compartment specified by duration parameter
14. First order bolus into central compartment at ss followed by an ss=2 (superposition ss) dose at 12 hours
15. First order bolus into central compartment at ss followed by an ss=2 (superposition ss) dose at 12 hours followed by reset ss=1 dose at 24 hours
16. Two parallel first order absorption models
17. Mixed zero and first order absorption

## Maximum Likelihood Tests

The Pumas-NLME parameter estimation algorithms, specifically *FOCE(), LaplaceI()*, and *SAEM()*, were compared with NONMEM in a richly-sampled data setting with multiple models. After a single administration 19 samples were collected over 72 hours via simulations for 4 different dose levels of 30 subjects each. A total of 6 test cases were generated that were a combination of:

- one- or two-compartmental disposition [26]
- oral (first-order absorption), intravenous (IV) bolus, or IV infusion administration.

The *Ω*’s, representing the between subject variability were set to 30% CV, implemented as diagonal covariance matrix. A proportional residual error with 20% CV was chosen to capture the error distribution. For the structural model, all one-compartment models had a population *V c* of 70 L, and all two-compartment models had an additional peripheral volume (*V*_*p*_) of 40 L. For all oral absorption models, *K a* was set to 1.0 h-1. All models with linear elimination had a *CL* of 4.0 L/h. All two-compartment models had inter-compartmental clearance (*Q*) set to 4.0 L/h.

## TMDD Model Tests

While the maximum likelihood tests above compared the standard compartmental models, we also evaluted the performance of highly non-linear mechanistic models in the class of Target Mediated Drug Disposition (TMDD). Three models from this class were selected:

1. Michaelis-Menten (MM) model
2. Constant Rtot
3. Rapid binding (QE) and quasi steady-state (QSS) models

Data were simulated from a Full TMDD model in 224 subjects over a range of doses with a total of 3948 observations. The simulated data were then estimated using the reduced models listed above using NONMEM and Pumas

## NCA Implementation Verification

Simulations were performed in R with scenarios consisting of 1-, 2-, and 3-compartment models with typical parameters and ±4-fold from those typical parameters on the ratio of absorption rate (*K a*) to elimination rate (*Kel*), ratio of peripheral volume of distribution 1 and 2 (*V p*1 and *V p*2) to central volume of distribution (*V c*), intercompartmental clearance between *V c* and *V p*1 or *V p*2 (*Qcp*1 and *Qcp*2), and ratio of clearance (*C*L) to *V c* all models, as the parameters apply; with and without target-mediated drug disposition (TMDD); and oral and intravascular bolus dosing. All models were simulated with 4%, 10%, and 20% proportional residual error. Each model was simulated with 6 subjects. This yielded a total of 13104 scenarios and 78624 subjects simulated.

Each subject was then grouped with all other subjects in its simulation scenario, and 5, 10, and 20% of concentration measurements were set to below the limit of quantification (LOQ). NCA was to be performed on each of those LOQ scenarios in PKNCA [9], Pumas, and Phoenix^4^ (a total of 707616 NCA intervals with calculations). Comparisons were made between the results of those NCA calculations performed on a single machine.

Five parameters, were chosen as the metrics for comparison as they included both observed and derived parameters:

1. AUClast
2. Cmax
3. Tmax
4. Half-life
5. AUCinf(pred)

Eighteen percent of subjects were randomly selected from the 13104 scenarios to form a subset of 2367 subjects. This smaller subset was used to perform the NCA calculations across the three software while ensuring to maintain the same default options. All analysis were conducted on the same laptop and discussed at ACoP 2019 [50].

Results from all three software for the key NCA parameters match. Most results matched within ±0.1% between all software. The difference between PKNCA /Pumas and Phoenix in half-life appears to be the result of PKNCA/Pumas selecting the best fit first and then filtering for decreasing slope while Phoenix first considers only consecutive sets of points that generate a descending slope and then selects the final set of points with the best regression adjusted R squared. None of the different results would be reported in a typical reporting workflow as all *r*^2^ values were <0.7.

## Validation Figures and Tables

The following plots verify the accuracy against widely used software for the computational experiments and benchmarks.

**Figure 15:**
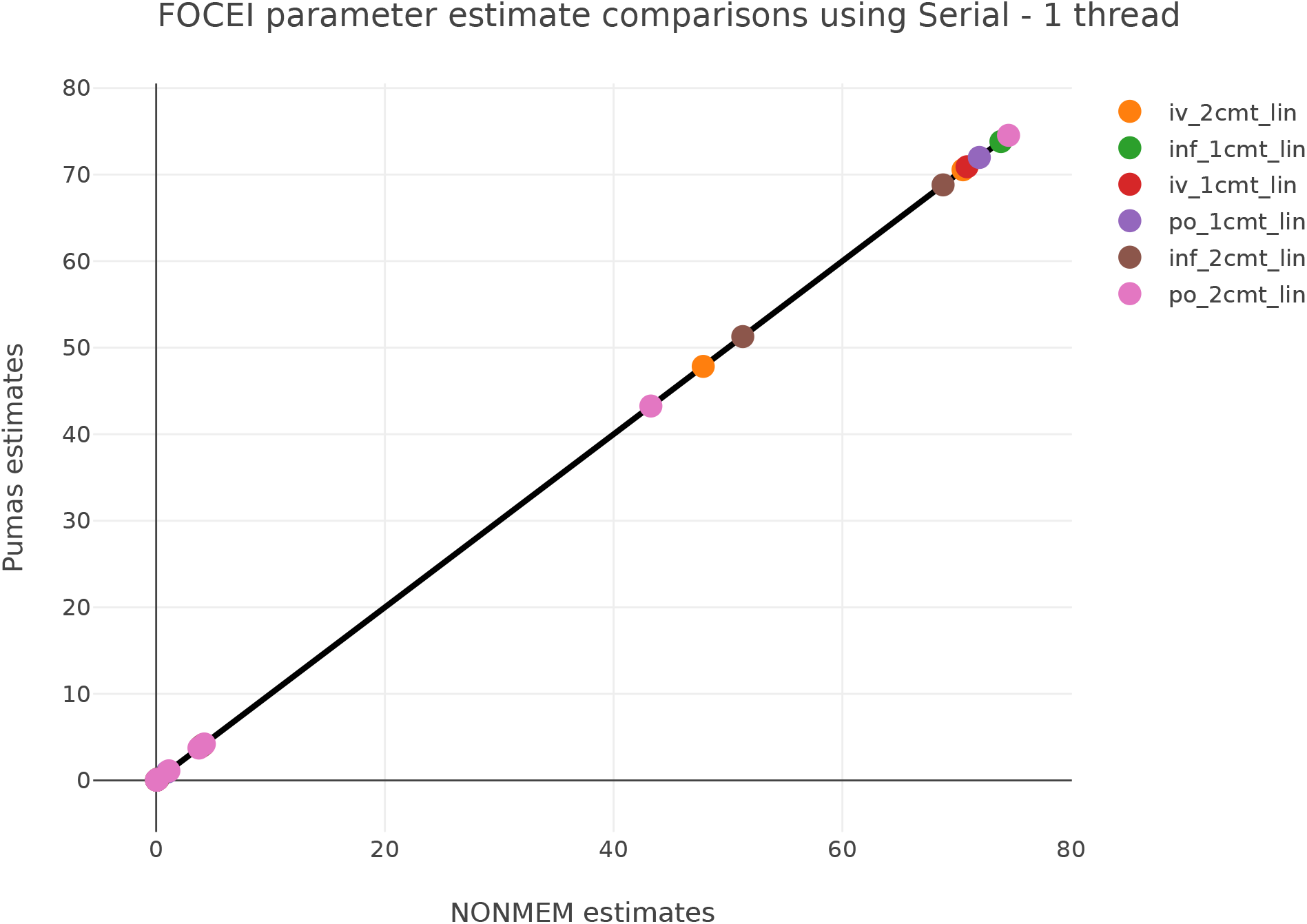
FOCE Benchmark Estimates. Shown are the resulting estimated values of the benchmarks in Figure 6 comparing the results of Pumas and NONMEM. Each point corresponds to one parameter estimated in the given benchmark. The *x* = *y* line indicates that the estimated values between the two software are the same.

**Figure 16:**
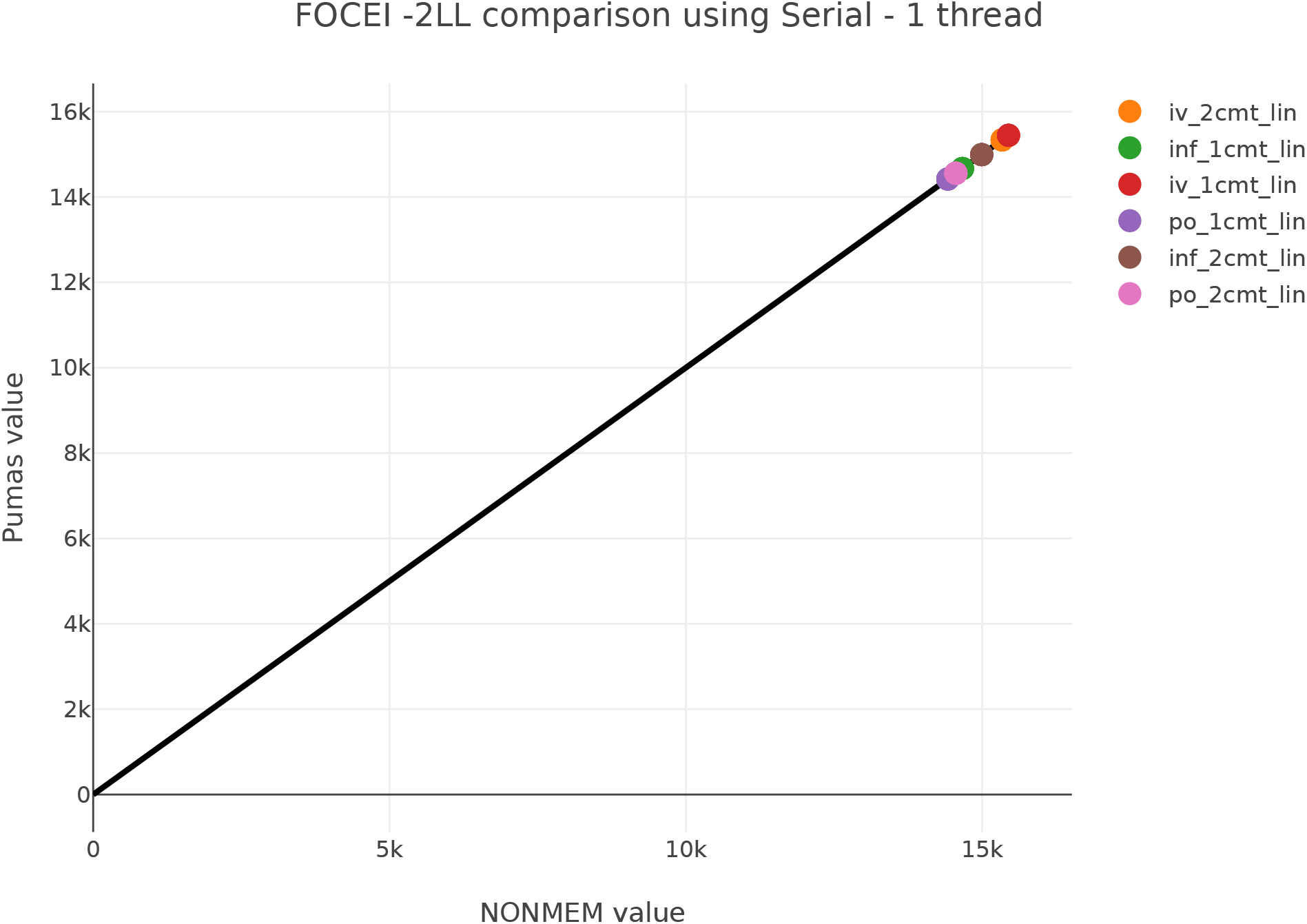
FOCE Benchmark Log-likelihood Results. Shown are the resulting estimated values −2loglikelihood fit results of the benchmarks in Figure 6 comparing the results of Pumas and NONMEM. Each point corresponds to the final log-likelihood of the given benchmark. The *x* = *y* line indicates that the estimated values between the two software are the same.

**Figure 17:**
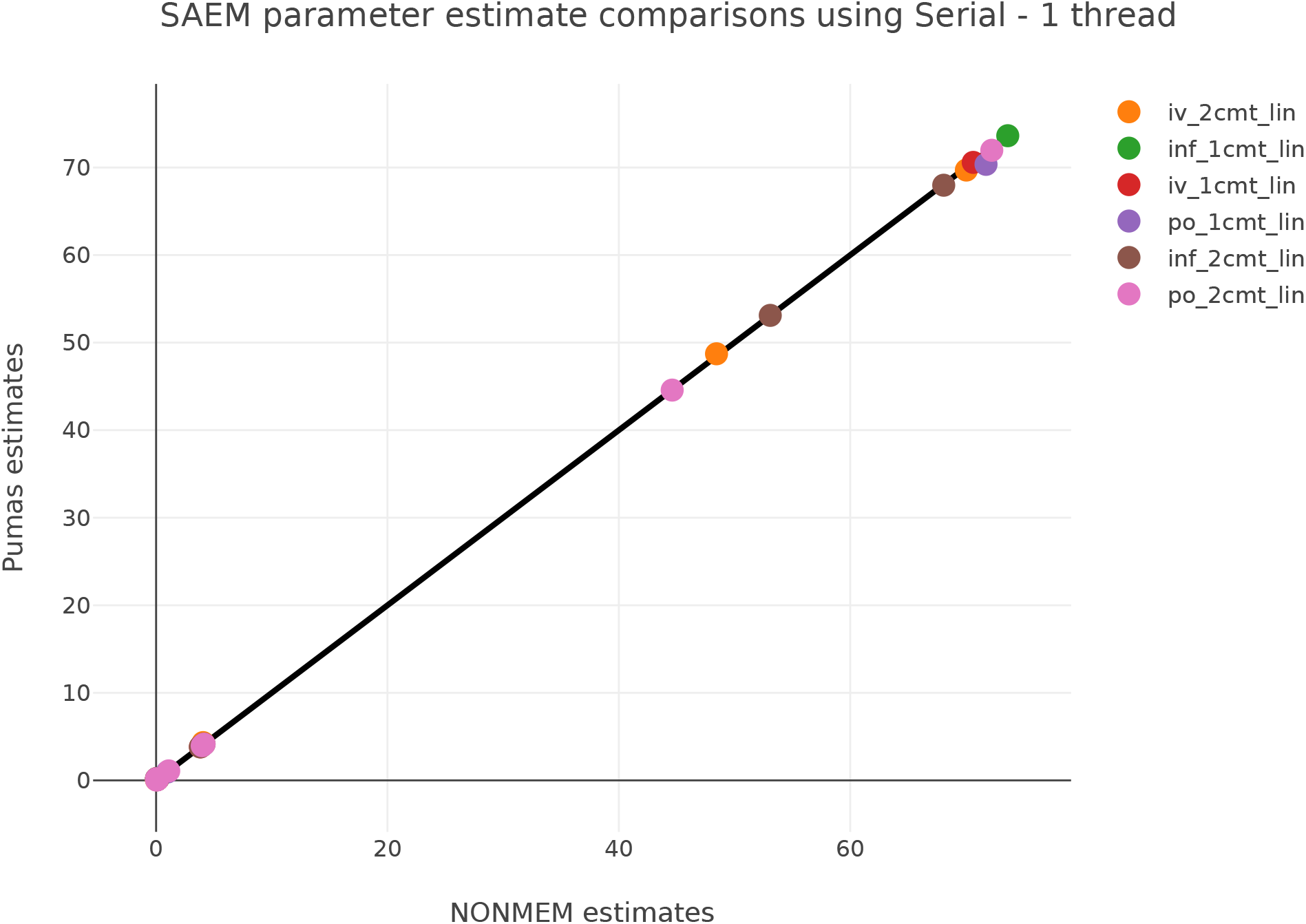
SAEM Benchmark Estimates. Shown are the resulting estimated values of the benchmarks in Figure 6 comparing the results of Pumas and NONMEM. Each point corresponds to one parameter estimated in the benchmark. The *x* = *y* line indicates that the estimated values between the two software are the same.

## Footnotes

1 Prior distributions are required for this fitting method and are specified in the `param block by using a distribution instead of a domain`.

2 defined in [28] unless mentioned otherwise

3 https://github.com/mrgsolve/nmtests/blob/master/nmtest7.md

4 https://www.certara.com/software/phoenix-nlme/

